# Multi-taxon inventory reveals highly consistent biodiversity responses to ecospace variation

**DOI:** 10.1101/807321

**Authors:** Ane Kirstine Brunbjerg, Hans Henrik Bruun, Lars Dalby, Aimée T. Classen, Camilla Fløjgaard, Tobias G. Frøslev, Oskar Liset Pryds Hansen, Toke Thomas Høye, Jesper Erenskjold Moeslund, Jens-Christian Svenning, Rasmus Ejrnæs

**Affiliations:** Department of Bioscience, Aarhus University, 8410 Rønde, Denmark; Department of Biology, University of Copenhagen, 2100 Copenhagen, Denmark; Rubenstein School of Environment and Natural Resources, The University of Vermont, Burlington, Vermont, USA; The Gund Institute for Environment, University of Vermont, Burlington, Vermont, USA; Section for Ecoinformatics and Biodiversity, Department of Bioscience, Aarhus University, 8000 Aarhus C, Denmark; Center for Biodiversity Dynamics in a Changing World (BIOCHANGE), Department of Bioscience, 8000 Aarhus C, Denmark; Natural History Museum Aarhus, Wilhelm Meyers Allé 10, 8000 Aarhuc C; Arctic Research Centre, Aarhus University, 8000 Aarhus C, Denmark

**Keywords:** abiotic environment, carbon resources, environmental DNA, environmental gradients, heterotrophs, intermediate productivity, primary producers, species richness, taxonomic aggregation

## Abstract

Amidst the global biodiversity crisis, identifying drivers of biodiversity variation remains a key challenge. Scientific consensus is limited to a few macroecological rules, such as species richness increasing with area, which provide limited guidance for conservation. In fact, few agreed ecological principles apply at the scale of sites or reserve management, partly because most community-level studies are restricted to single habitat types and species groups. We used the recently proposed *ecospace* framework and a comprehensive data set for aggregating environmental variation to predict multi-taxon diversity. We studied richness of plants, fungi, and arthropods in 130 sites representing the major terrestrial habitat types in Denmark. We found the abiotic environment (ecospace position) to be pivotal for the richness of primary producers (vascular plants, mosses, and lichens) and, more surprisingly, little support for ecospace continuity as a driver. A peak in richness at intermediate productivity adds new empirical evidence to a long-standing debate over biodiversity responses to productivity. Finally, we discovered a dominant and positive response of fungi and insect richness to organic matter accumulation and diversification (ecospace expansion). Two simple models of producer and consumer richness accounted for 77 % of the variation in multi-taxon species richness suggesting a significant potential for generalization beyond individual species responses. Our study widens the traditional conservation focus on vegetation and vertebrate populations unravelling the importance of diversification of carbon resources for diverse heterotrophs, such as fungi and insects.

## Introduction

For centuries, ecologists have struggled to understand and explain spatial and temporal variation in biodiversity, with increasing societal attention motivated by the global biodiversity crisis. Most models and theories of biodiversity refer to specific taxonomic groups and ecosystems (1–6), leaving us with no general rules and ecological theories of diversity and without prioritization tools at the local spatial scales of practical conservation planning and management. Several studies have investigated the potential for using selected species groups to represent the wider conservation interests, but based on a global meta-analysis, Westgate, Barton, Lane and Lindenmayer (7) concluded that the data undermines the assumption that a taxonomic subset can represent the wider biodiversity. The disappointing conclusion has been somewhat contradicted or modified lately by studies showing promising potential for cross-taxon congruence in species composition and compositional turnover (β-diversity) (8) and also for species richness, but only after accounting for environmental variation (3, 9). Along this line of reasoning, we set out to test the hypothesis that terrestrial multi-taxon diversity can be predicted across contrasting environments without detailed consideration of an intractable diversity of taxonomic groups, response groups or habitat types. We are thus not testing the surrogacy hypothesis per se, but rather the idea that multi-taxon diversity can be predicted from a low-dimensional ecological space, ignoring the possible multitude of response shapes of the individual taxonomic groups or species.

We applied the recently proposed *ecospace* framework (2–4, 10, 11) for a formal and structured quantification of environmental variation. Ecospace represents the total environment in space and time, in which the individual (species) colonizes, grows, reproduces, and dies or goes extinct. Ecospace may help reduce environmental complexity to a tractable number of dimensions and measurable variables. We have proposed to subdivide ecospace into three components each signifying important aspects of an area for its potential biota: 1) The abiotic environment (*ecospace position*), 2) The accumulation and diversification of organic matter (*ecospace expansion*) and 3) The spatio-temporal *continuity* (10).

The position of a site in n-dimensional environmental hyperspace (e.g., mean values of soil moisture, pH, soil fertility, and temperature) is essential to sessile organisms like plants and soil-fungi unable to move across their local environment (6), but even mobile animals respond to abiotic conditions when they select their habitat (12). Expansion, i.e. the accumulation and diversification of organic matter, is particularly important as an energy source for consumers (13), but may also provide substrate for e.g. epiphytic plants and lichens (14). Expansion presupposes primary production and subsequent differentiation into leaves, roots, stems, flowers, bark, wood, dung etc. On evolutionary time scales, every differentiated pool of live or dead organic matter has provided opportunity for heterotrophic niche differentiation and speciation (15).

The spatial continuity of habitats is expected to be particularly important for short-lived and poorly-dispersed species moving among patches varying in suitability over time, but less important for species with long distance dispersal (16). Temporal continuity on the other hand should be particularly important for organisms with limited dispersal ability and high persistence, such as some plants and fungi, which are sensitive to changing habitat conditions (6).

The aim of this study was to investigate the hypothesis that multi-taxon α-diversity, including ‘genetic richness’ (5) from environmental DNA, can be treated as a general, predictable biotic response to environmental variation represented by a low number of key factors. We assess this using ecospace as a framework for guiding the study design, environmental mapping and data analysis.

## Results

Individually, the nine species richness models explained between 27 % (flying insects) and 62 % (decomposing fungi, herbivorous arthropods) of the variance in species richness across environmental gradients (Fig. 2). When we pooled species into producers and consumers, the explained variation based on cross-validated predictions was comparable to the individual models (producers: 49 %, predictions producers: 50 %; consumers: 73 %, predictions consumers: 69 %). The model for species richness of all groups explained 54 % of the variation in biodiversity across sites, considerably less than the predictions from the sum of all nine separate species group models (74 %). However, the summed predictions from the models of producers and consumers explained 77 % of total species richness. Based on this result, we focused further reporting of results on models of producer and consumer biodiversity (Fig. 3). Producer richness was primarily explained by position (Fig. 3a); increasing with soil pH, intermediate nutrient status, extreme soil moisture (wet/dry sites), presence of boulders and with the ecological plant species pool size. The ecological species pool index (index based on vascular plants that reflect the importance of evolutionary and historical contingency on local community assembly) reveals that there are more vascular plants in the Danish flora which prefer relatively high incoming light, intermediate soil moisture and relatively high soil pH (Fig. S1). Presence of a shrub layer also promoted producer richness. Finally, producer richness increased with temporal continuity. Variance partitioning revealed that position explained most variation in producer richness with minor contributions from expansion and continuity (Fig. 3a). Consumer richness increased with presence of a shrub layer, higher flower abundance, and a high index of insect host plant abundance (Fig. 3b). For position, consumer richness increased with air temperature and decreased with incoming light. We found no effect of continuity on consumer richness. Variance partitioning revealed that most variation in consumer richness could be explained by expansion compared to a minor contribution from position (Fig. 3b).

**Figure 1:**
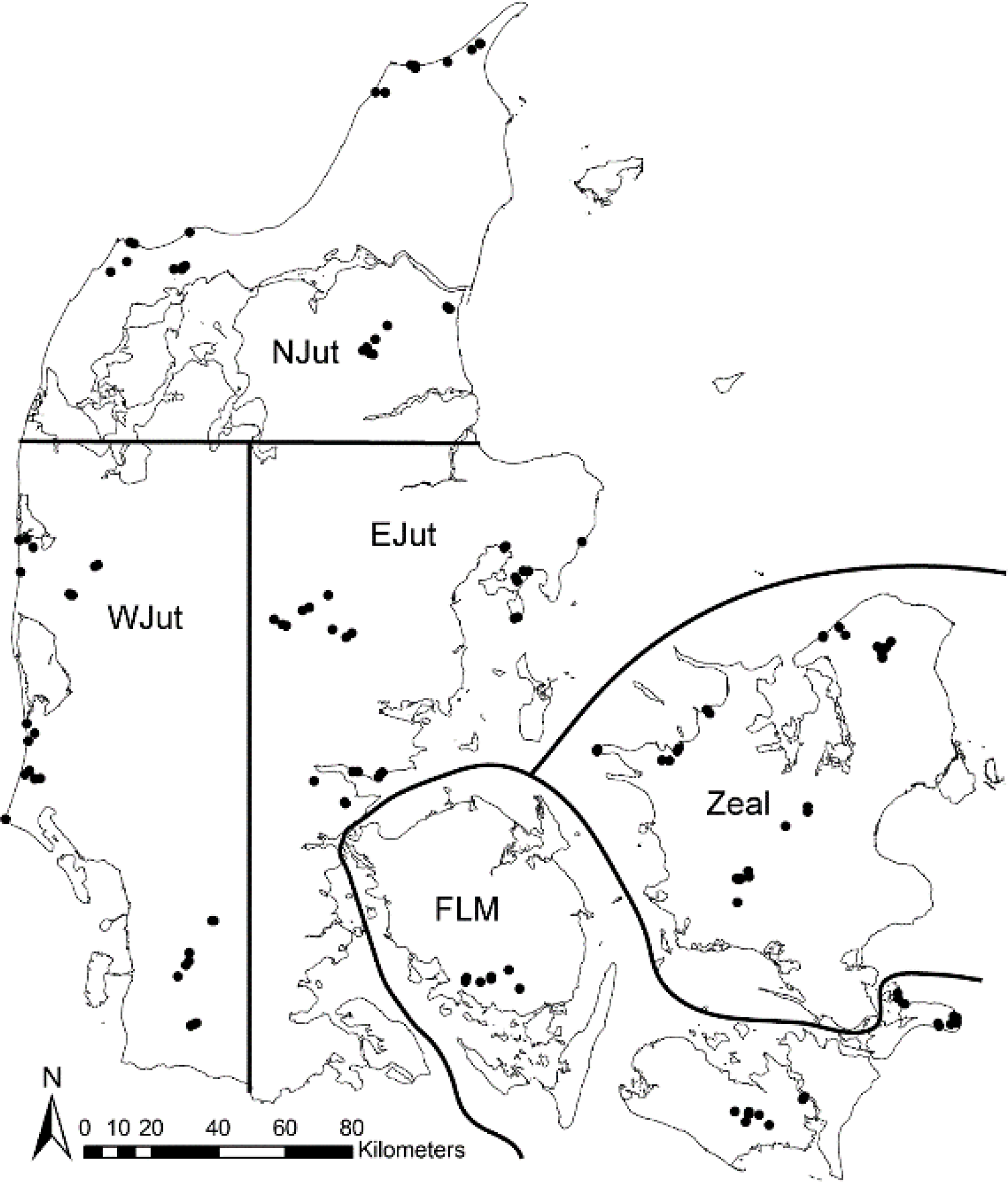
Map of Denmark showing the location of the 130 sites grouped into 15 clusters within five regions. (NJut: Northern Jutland; WJut: Western Jutland; EJut: Eastern Jutland; FLM: Funen, Lolland, Møn; Zeal: Zealand). Reprinted from Biological Conservation, 225, Ejrnæs R, Frøslev TG, Høye TT, Kjøller R, Oddershede A, Brunbjerg AK, Hansen AJ, & Bruun HH, Uniquity: A general metric for biotic uniqueness of sites, 98-105, Copyright (2018), with permission from Elsevier

**Figure 2:**
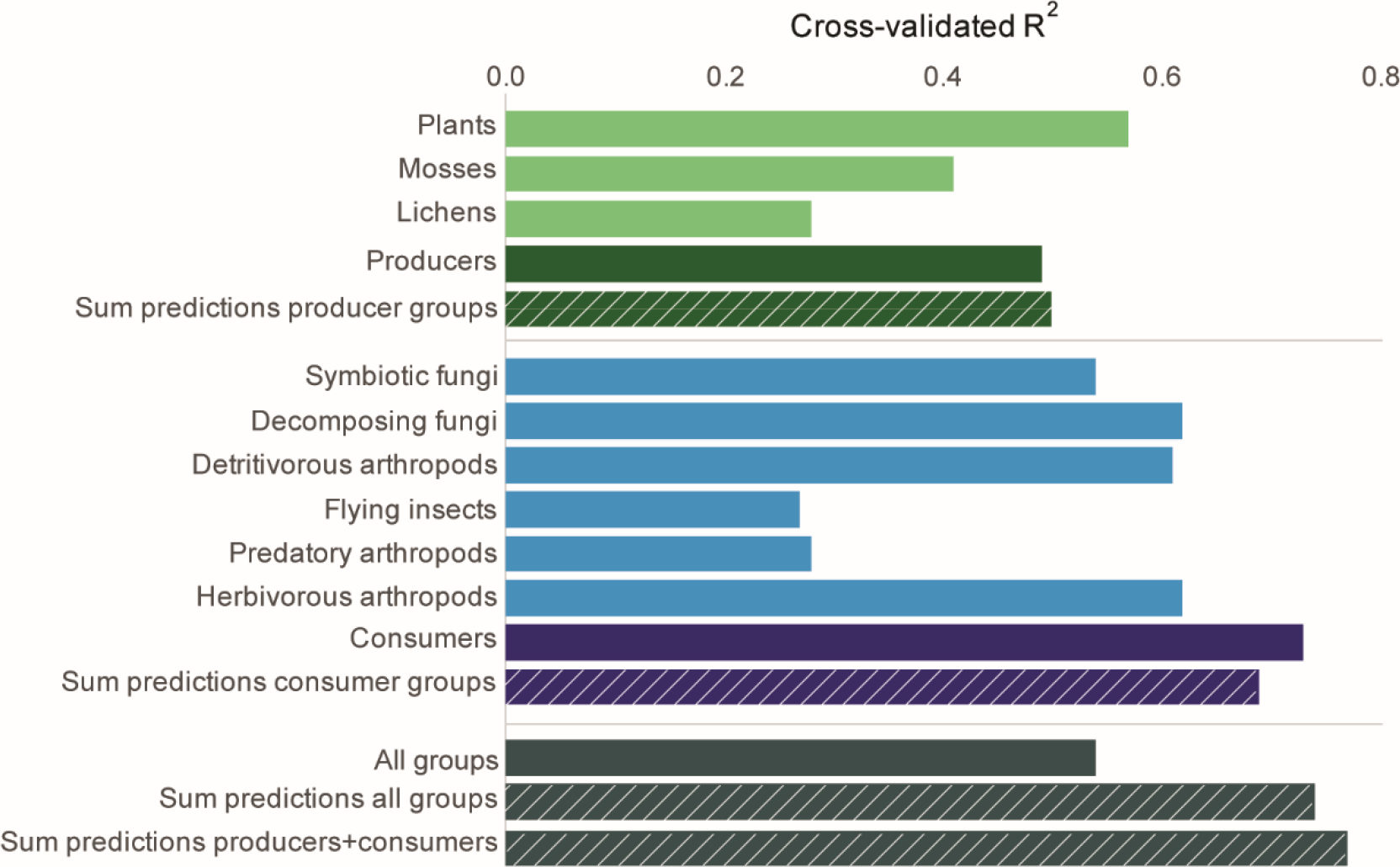
Cross-validated R^2^-values (%) for the best GLM negative binomial models explaining species richness in 130 sites from environmental and geographic variables. Green – Producers, blue – Consumers, dark grey – All groups species richness. Hatched bars represent explained variation from pooling the predictions of individual models and they are included for evaluating the effect of taxonomic aggregation to the level of producers, consumers and all groups species richness. For example, the hatched dark green bar is the correlation between the pooled predictions from the plant, moss and lichen models respectively and the observed richness for all of these groups. This compares to the solid dark green bar representing the correlation between the predictions from a model including all producer groups and the observed values. For all groups, the first hatched bar represents the correlation for the summed predictions from all nine species group models, whereas the second hatched bar represents the correlation for the summed predictions for the producer and the consumer models.

**Figure 3:**
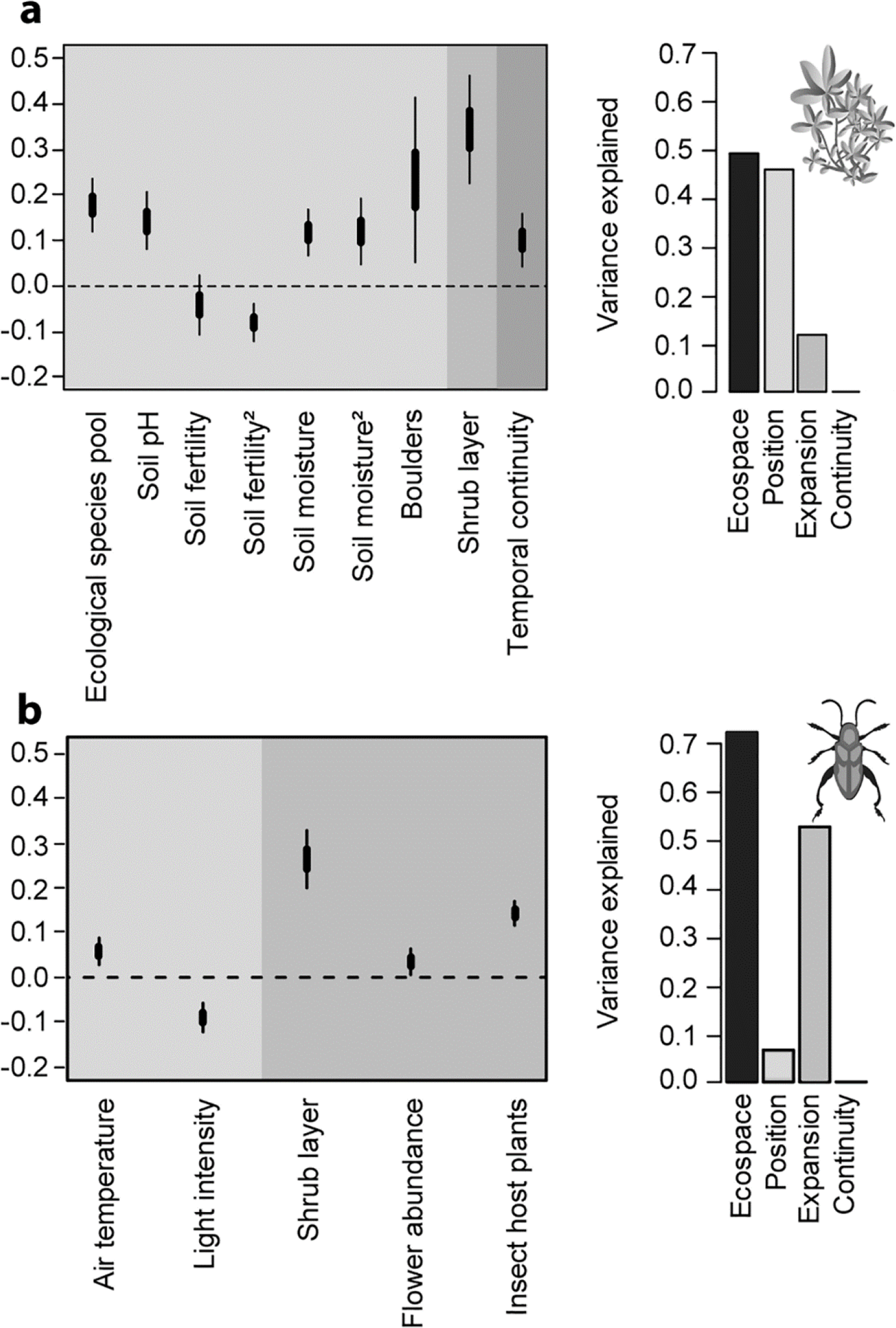
Coefficient plot of the best models for a) producer (plants, mosses, lichens) and b) consumer (fungi, arthropods) richness in the 130 sites (left-hand panel) and explained variance for ecospace and its components (position, expansion, continuity; in the right-hand panel). Explanatory variables are standardized and shaded according to ecospace components: position – light grey, expansion – grey, continuity - dark grey. Thick (inner) bars represent ±1 standard error, thin (outer) bars represent ±2 standard errors.

The following expansion variables promoted species richness for the individual response groups: litter mass for decomposing fungi, soil carbon content for arthropod detritivores, dung for decomposing and symbiotic fungi as well as total species richness, dead wood for decomposing fungi and floral abundance for total species richness, consumers, flying insects and herbivorous arthropods (Table 1). Richness of all consumer species groups increased with either plant species richness or the indices of host plant availability for fungi or insects indicating a bottom-up effect going from primary producer richness to consumer richness. The presence of a shrub layer had a consistent and positive effect on the richness of most response groups. We found very few significant effects of within-site heterogeneity on species richness indicating that these were of little importance compared to the effects of between-site variability (Table 1).

**Table 1:**
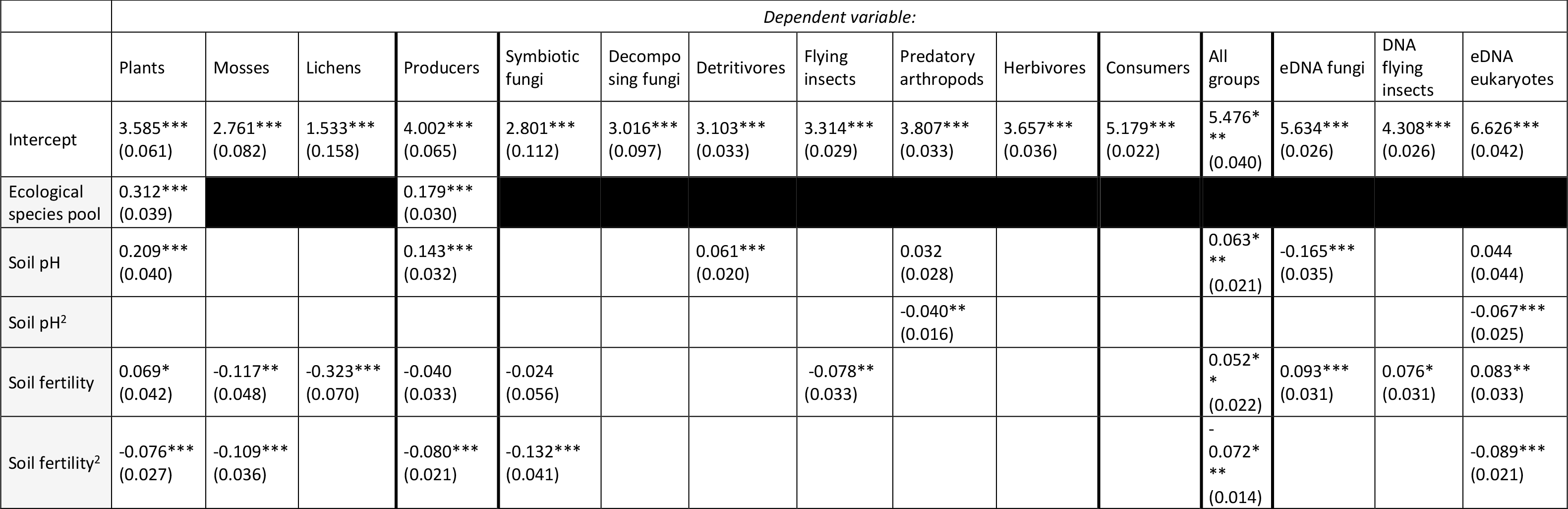
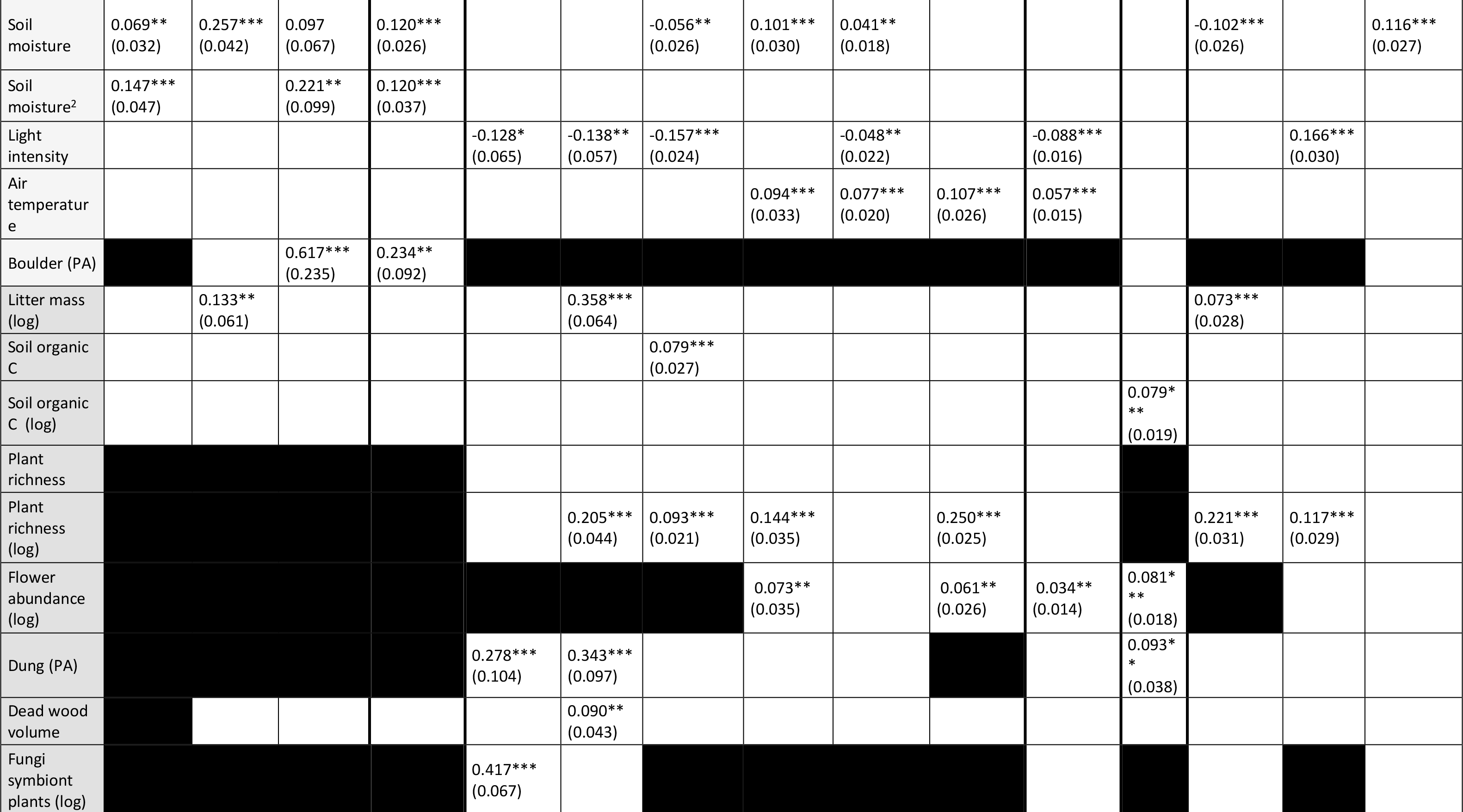
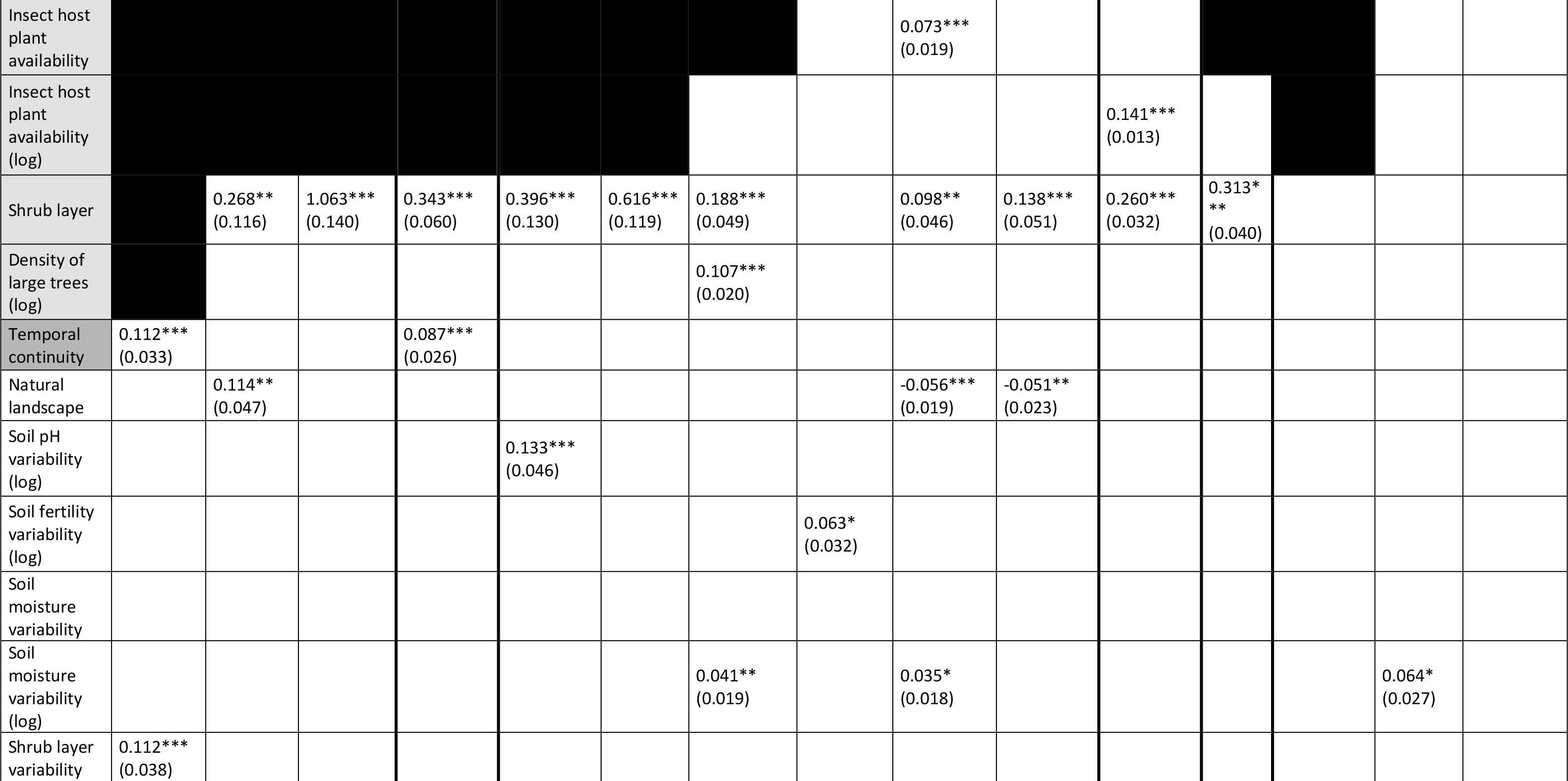
Model output for GLM negative binomial models using site (n=130) richness of plants, mosses, lichens, producers (plants, mosses and lichens), symbiotic fungi, decomposing fungi, detritivores (poisson), flying insects, predatory arthropods, herbivores, consumers (fungi and arthropods), all groups, eDNA fungi, eDNA eukaryotes, DNA flying insects. Estimates, p-values (* <0.05, ** <0.01, *** <0.001) and standard errors (in parentheses) are given. Explanatory variables are colored according to the ecospace framework: position – light grey, expansion – grey, continuity – dark grey, co-variables – white. Black: variable not relevant for species group (see Supplementary Information Table S4).

In general, we found linear richness responses to underlying abiotic gradients, with the exception of unimodal responses of predatory arthropods to pH, and bimodal responses of lichens, vascular plants and producers to soil moisture. For soil fertility, we observed unimodal responses for vascular plants, mosses, symbiotic fungi as well as the pooled groups of producers and total species richness (Fig. 4).

**Figure 4:**
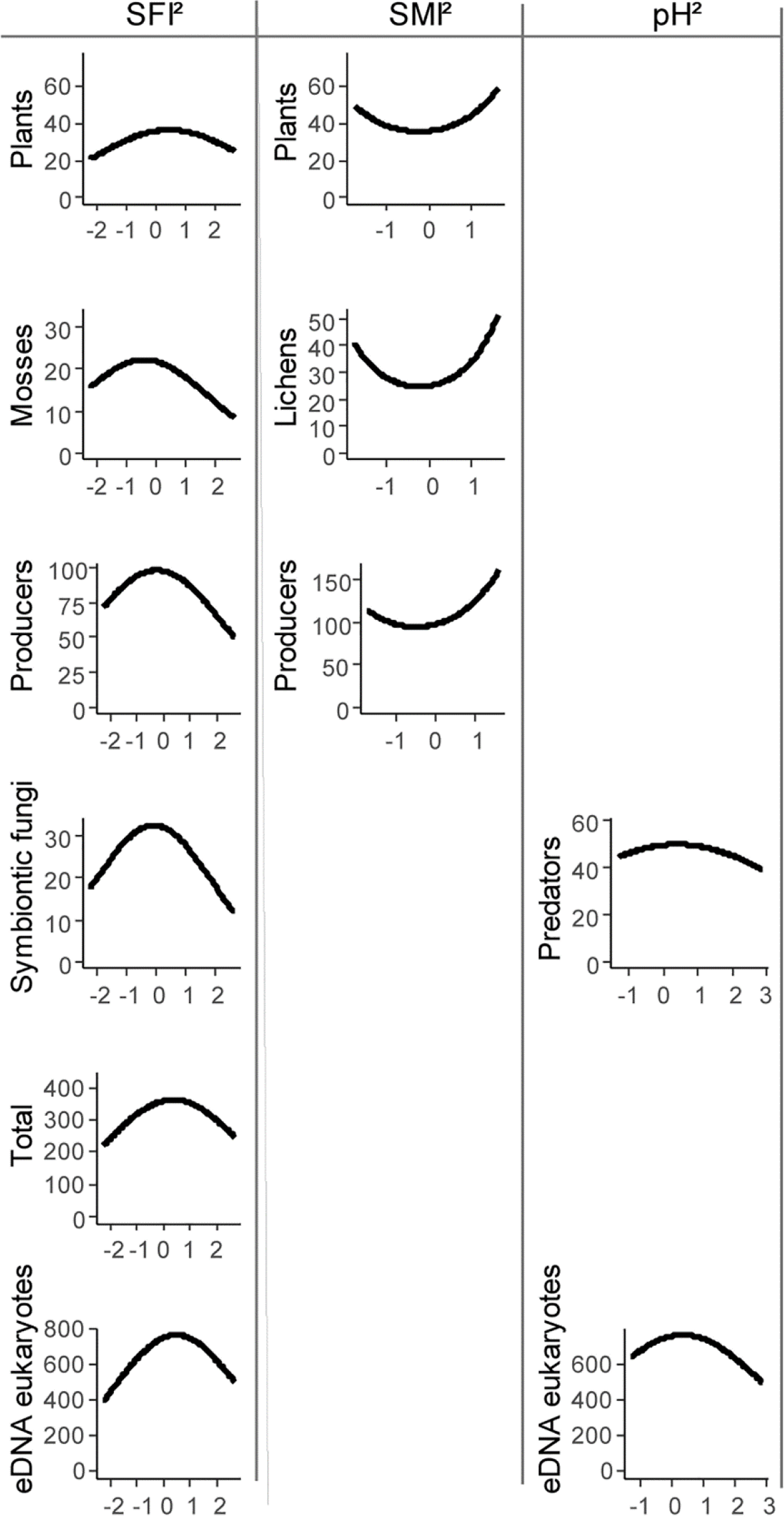
Relationships between significant squared ecospace position terms for soil fertility (SFI), soil moisture (SMI) and pH and the species richness of plants, mosses, lichens, producers, symbiotic fungi, predatory arthropods, total (producers+consumers), and eDNA eukaryotes in the respective multiple regression models.

Models for ‘genetic richness’ of soil and insect trap DNA were in the lower range of model performance with cross-validated R^2^ values of 23 % for eukaryotes, 27 % for fungi and 29 % for flying insects. While fungi and flying insects confirmed the consistent positive response to plant richness, none of the DNA groups responded to presence of a shrub layer (Table 1). Soil pH, moisture, fertility and litter mass affected soil fungal and eukaryotic DNA richness in various ways indicating the importance of the soil environment for its biota (Table 1).

We found no justification for including a random variable (Generalized Linear Mixed Model) to account for spatial patterns in the data and only two of 15 models (the flying insects and DNA flying insects models) showed significant spatial autocorrelation after modelling. Spatial signals seem to be of minor importance in this study.

## Discussion

Across major terrestrial ecosystems within a region, as much as 77 % of the variation in multi-taxon richness was accounted for by our models. The remarkably high explanatory power arose after grouping species into producers (autotrophic organisms) and consumers (heterotrophic organisms). Studies assessing the surrogacy power of multiple environmental variables on multiple taxonomic groups simultaneously, are rare. A meta-study found the surrogacy power of environmental variables to be very poor compared to taxonomic surrogacy (17) while multi-metric site-conditions explained 48 % of variation in total species richness of a range of taxa (ants, beetles, spiders, wasps, flies, butterflies, reptiles, birds, vascular plants, bryophytes and lichens) in Australian woodlands (18). Our regression models for individual species groups show considerable variation in the selection of variables and this could easily lead to the erroneous conclusion that each taxonomic group needs individual consideration. Nevertheless, the predictions derived from models dedicated to taxonomically and ecologically more specialized species groups explained less variation in α-diversity than the two models based on high level aggregation. This level of generalization is striking considering the contrasting life history traits and modes of resource acquisition of the species aggregated as producers or consumers, and it challenges commonly expressed concerns that α-diversity is necessarily contingent on taxonomy and ecology (e.g., 19, 20, 21).

On the other hand, we found a large drop in explained variation when we tried to aggregate producers and consumers and model all species in one model. This result emphasizes the fundamental difference between sessile autotrophic organisms, whose resource acquisition is largely controlled by abiotic conditions, and heterotrophic organisms, whose resource acquisition relies on biotic resources. The important split between heterotrophic and autotrophic organisms is underpinned by the contrasting role of ecospace position and expansion in the producer and consumer models. Further, plant richness is an important predictor of consumer richness, but to avoid circularity plant richness is excluded from the total richness model.

Based on island biogeography (14) and metapopulation theory (15) we would have expected that ecospaces at larger spatial and temporal extent would increase the probability of immigration and decrease the risk of extinction. Surprisingly, spatio-temporal continuity played a negligible role in explaining multitaxon species richness. Only temporal continuity appeared as a small significant effect in the model of vascular plants and in the producer model. This result does not preclude continuity playing an important role in other biomes or geographical contexts (22). The species pool index for vascular plants emerged with a significant positive effect in the models of vascular plants and producers. This supports the biogeographic theory that species pools founded in evolutionary and historical timescales have lasting impacts on current biodiversity (23). The consistent positive effects of vascular plant richness and host plant indices on fungal and insect richness (Fig 3b, Table 1) corroborates a similar species pool effect for consumers (24). Despite recent advances in the translation of eDNA data into diversity metrics (25, 26) the relatively poor performance of models for DNA groups might point to remaining metagenomic challenges (25, 27). It is also possible however that the variation in below-ground and above-ground biodiversity is determined by different factors and that ecospace expansion in particular may need to be refined to include and differentiate between organic matter pools that support below-ground biodiversity (28).

The most consistent predictors of consumer richness were the presence of a shrub layer and high vascular plant richness. Specialized organic matter, such as dead wood, dung and flowers were found to be important to fungal and insect richness (29–31), judged from their representation in the detailed species group models. In this way, our study reveals a bottom up regulation of species richness emphasizing the importance in nature management of the build up of growing plants and the differentiation of species richness in the vegetation. Our result also implies that conservation managers should ensure effective protection against harvesting and homogenizing of organic matter such as live vegetation, flowers, and dead wood. Our results support the notion of large herbivores as keystone species promoting local richness by suppression of dominant plants (32), as long as the grazing regime does not obstruct the annual build up and flowering of the herb layer as well as the long-term build up of complex vegetation including a shrub layer, veteran trees and dead wood. The provision of large dung and the occasional damage to live trees should be considered instrumental to the diversification of organic matter. We envision that the role of natural dynamic processes for the diversification of organic matter – not least in the soil (33) – will be a promising field of research in future conservation studies.

To mitigate the biodiversity crisis, there is a need for ecological rules and principles to inform conservation planning and restoration actions (34). Without disregarding the scale dependence of biodiversity (35, 36) and the importance of endemism and threatened species (37), our study has demonstrated unprecedented potential for generalization of multi-taxon species richness responses to environmental variation, supporting our hypothesis of α-diversity as a predictable property of a low-dimensional ecospace.

## Methods

During 2014-2017, we collected data from 130 sites (40 m × 40 m) within 15 clusters nested in five regions across Denmark (Fig. 1). We allocated 100 sites to span the most important natural gradients affecting biodiversity in Denmark i.e. gradients in soil fertility, soil moisture and successional stage: from nutrient rich to nutrient poor, from dry over moist to wet and from open, closed and forested vegetation. 90 of these sites were selected randomly within 18 predefined strata, whereas 10 sites were selected by the amateur natural historian community to represent biodiversity hotspots for different species groups. The remaining 30 sites were sampled randomly from six strata cultivated for production: plantations (beech, oak or spruce) and fields (rotational, grass leys or set aside). Randomization was achieved by selection of sampling areas and potential sites from the office desk based on geo-referenced information – for some rare strata we also consulted local experts. To minimize spatial autocorrelation, the minimum distance among sites was 500 m with a mean nearest distance among sites of 2291 m. Within each site, we sampled vascular plants, mosses lichens, fungi, and arthropods, and we included metabarcoding of DNA from soil samples and insect traps to reflect the ‘genetic richness’ (i.e. number of operational taxonomic units – OTUs (5) of eukaryotes, fungi and flying insects.

Furthermore, we collected data reflecting ecospace i.e. abiotic position, biotic expansion and spatio-temporal continuity (described below). For further details on site selection and data collection, see Brunbjerg, *et al.* (11). All field work and sampling was conducted in accordance with Responsible Research at Aarhus University and Danish law.

### Ecospace position variables

ecospace position represents the abiotic factors affecting species occurrence directly via environmental filtering (10) or indirectly causing variation in species pools developed over evolutionary and historical temporal scales. The following variables represented abiotic position:

#### Ecological species pool index

We developed the ecological species pool index to reflect the importance of evolutionary and historical contingency on local community assembly. The species pool was only developed for vascular plants as we did not have access to independent data for other species groups (but see host plant indices for fungi and insects below). We extracted Ellenberg Indicator Values (38) for all vascular plants considered part of the Danish flora (www.allearter.dk). Ellenberg Indicator Values specifies plant optima for ecological conditions and we used light (Ellenberg L), moisture (Ellenberg F) and pH (Ellenberg R) as predictors of species pool size. We avoided Ellenberg nutrient preference, as this indicator implies competitive hierarchies. We used a quasi-poisson GAM-model (k=3) with Ellenberg values as explanatory variables and the number of plant species associated with each unique combination of Ellenberg F, L and R as a response variable (the three variables were normalized to 0-1 before modelling). To estimate the species pool index for each of the 130 sites we predicted the number of plants based on the GAM-model and the site mean Ellenberg F, L and R values. The species pool index was log-transformed as this provided the best linear fit to observed species richness.

#### Soil Fertility Index (SFI)

Soil fertility is a complex attribute involving cycling, holding capacity, release rate, immobilization, leaching etc. and nutrient availability changes over the season. We chose to calculate a soil fertility index by integrating a range of abiotic variable measures/indicators of nutrient content or cycling. For each site SFI represents the predicted value from the best linear model (of all sites) of site mean Ellenberg N (38) (plant-based bioindication of nutrient status) as a function of soil calcium (Ca), leaf nitrogen (N), leaf N:phosphorus (NP) and soil type (Table S2).

#### Soil Moisture Index (SMI)

Soil moisture is a complex attribute reflecting local hydrology as well as precipitation and soil moisture varies between seasons and years. We calculated a soil moisture index for each site using the predicted values from the best linear model (of all sites) of mean Ellenberg F (38) (plant-based bioindication of soil moisture) as a function of mean precipitation in 2001-2010 (10 km × 10 km grid resolution) and measured site soil moisture (trimmed mean of 16 measures per site taken with a FieldScout TDR 300 Soil Moisture Meter in May 2016).

#### Soil pH

We measured soil pH on soil bulk samples (mix of four samples per site: depth = 0-10 cm, 5 cm diameter).

#### Light

We measured light intensity (lux) reflected microclimate in each site using HOBO Pendant^®^ Temperature/Light 8K Data Loggers.

#### Air temperature

Air temperature (°C) reflected microclimate in the sites and was measured using HOBO U23 Pro v2 Temperature/Relative Humidity data loggers.

#### Boulders

We measured the presence of boulders (diameter > 20 cm) within each site using presence-absence because there is a low density of boulders on the Danish landscape. While boulders are ‘habitats’ for epilithic mosses and lichens, here we defined them as position because of their abiotic nature.

### Ecospace expansion variables

Expansion is comprised of organic matter that species can live on (surfaces) or from (resources). Expansion variables reflect both quantitative (amount of organic matter) and qualitative (diversity of organic matter) aspects.

#### Pools of organic matter

1. Dung of herbivores (presence/absence of dung of hare, deer, sheep, cow or horse).
2. Litter mass (g/m^−2^ of four litter samples within a 21 cm × 21 cm frame per site).
3. Flowers: The density of flowers of insect-pollinated plants presenting flowers within the site (sum of estimates from June and August 2014 and April 2015). Flower density was recorded using plotless sampling (39) and weighted by flower surface area as follows: if flower surface area < 4 cm^2^ flower density was multiplied by 2, if flower surface area 4-10 cm^2^ flower density was multiplied by 7 and if flower surface area > 10 cm^2^ flower density was multiplied by 15.
4. Dead wood: diameter and length of coarse woody debris (>20 cm diameter, min length 1 m) was recorded and volume/ha was calculated.
5. Fine woody debris: density of fine woody debris (5-20 cm diameter and > 1 m, or >20 cm and < 1 m), including tree stumps within the site was recorded by plotless sampling (39).
6. Density of large trees: the number of large live trees (> 40 cm DBH) within the site was recorded.
7. Organic matter: percentage of the 0-10 cm soil core that was categorized as organic soil.
8. Soil organic C content: % soil C in 0-10 cm soil layer (g/m^2^ average of four soil samples taken in each site).
9. The number of plant species per site: as plants make up a carbon pool and structural habitats for fungi and arthropods (40) standardized number of plant species per site was used as expansion variable in fungi and arthropod models as well as eDNA eukaryote, eDNA fungi and flying insects models. While plant richness is a major predictor of consumer richness, it must be excluded from the total richness model in order to avoid circularity.

#### Shrub and tree layer

We subtracted the digital elevation model (DEM (41)) (40 cm × 40 cm resolution) from the digital surface model (40 cm × 40 cm resolution) to create a grid representing the above-ground vegetation height. From this, we calculated two variables for each site: The 90th percentile for returns > 3 m within the site reflecting the height of mainly trees (called tree layer) and the 90th percentile for returns 30 cm - 3 m reflecting the height of the shrub layer. The shrub layer was recalculated to a presence/absence variable splitting the data at 2 meters, motivated by a pronounced bimodal distribution of data points.

#### Indices for abundance of insect host plants and fungi host plants

In order to produce indices reflecting the availability of possible host plants for insects and fungi, we retrieved information on the associations between vascular plants and their consumers (fungi and insects). For fungi, we used observational data from the Danish Fungal Database (https://svampe.databasen.org) and for insects, we used accounts from an insect host plant database for NV Europe hosted at the Biological Records Centre (http://www.brc.ac.uk/dbif/hosts.aspx). Some consumers have links to many plant species while others are specialized to a single species or genus of plants. We assured equal importance of each consumer-link by weighting each link inversely with the number of plant genera involved with the consumer in question. Links reported to the genus level were attributed to all plant species within that genus. After summing all observed links for each plant species, we produced models predicting plant host attractiveness using plant functional traits as explanatory variables (unpublished work in progress). The model for fungi explained 66% of the observed link score and the model for insects explained 46 % of observed link score. We used the model to predict a value for each plant species in our data set, and we used the sum of values for the species of a site weighted by a species abundance score in the site (from 1-3) as index for host plant availability.

### Ecospace continuity variables

Continuity is the extent of the site habitat (position and expansion) in time and space.

#### Geographical species pool

The geographical species pool reflects the impact of historical processes on the species pool size under the assumption that immigration is ongoing and geographically directed (from Southeast towards Northwest) and that the Danish flora is not yet saturated. We estimated a geographical species pool for each site from predictions of a GAM model on vascular plant species richness as a function of geographic coordinates using an independent data set from a national Atlas Survey of vascular plant species in 1300 reference quadrats of 5 km × 5 km (42).

#### Spatial continuity

We estimated spatial continuity by assessing the amount (%) of natural areas within four different distances from the site (500 m, 1000 m, 2000 m and 5000 m). Spatial continuity of the habitat type of the site was estimated by visual interpretation of aerial photographs and additional information from land mapping of woodlands, fields, grassland, heathland, meadows, salt marshes and mires. The four buffer sizes were similar and highly correlated. The 500 m buffer was used for analyses as most of the studied species were expected to have relatively limited dispersal and small area requirements.

#### Temporal continuity

Temporal continuity was estimated by time since major land use change within the 40 m × 40 m site. For each site, a temporal sequence of aerial photos and historical maps was inspected starting with the most recent photos (photos from 2014, 2012, 2010, 2008, 2006, 2004, 2002, 1999, 1995, 1968, 1954, 1945) and ending with historical maps reflecting land use in the period 1842-1945. Temporal continuity (the year in which a change could be identified) was reclassified into a numeric 4-level variable: 1: 1-14 years, 2: 15-44 years, 3: 45-135 years, 4: >135 years.

### Co-variables

We include the following co-variables to account for a possible spillover of species by passive colonization from natural habitats in the surrounding landscape as well as a possible effect of within-site heterogeneity, increasing opportunities for niche differentiation. Co-variables were log transformed if transforming improved the distribution (visual inspection).

#### Natural landscape

We calculated the % share of natural or extensively used areas (forests, wetlands, heathlands and grassland) in 1 km × 1 km quadrats and interpolated these using Spline in Argis 10.2.2, Weight 0.5, number of points 9 (43).

#### Heterogeneity variables

##### Soil moisture variability

The variance of trimmed mean of 16 evenly distributed measurements of soil moisture within each site taken with a FieldScout TDR 300 Soil Moisture Meter in May 2016.

##### Soil fertility variability

The variance of four soil fertility index values per site (soil fertility index = predicted values of a linear model of Ellenberg N as a function of leaf N, leaf NP, soil P, soil Ca and soil class). Soil fertility variability was log transformed due to skewness.

##### Soil pH variability

The variance of four evenly distributed soil pH measurements per site.

##### Tree layer variability

The variance of the 90th percentile for returns > 3 m within the site reflecting the variability of the height of mainly trees (see description of the lidar high expansion variable above).

##### Shrub layer variability

The variance of the 90th percentile for returns 30 cm - 3 m reflecting the variability of the height of the shrub layer (see description of the lidar low expansion variable above).

#### Response variables

We divided species into response groups according to taxonomy (plants, mosses), trophic level (macrofungi) and trophic level and mobility (invertebrates). Grouping is complicated for insects because the biology and mobility may depend on live stage. Hoverflies for example have larval stages spanning from detritivores over predators to galling herbivores, whereas the adult flies mainly feed on flowers. An entirely trophic categorization is further intractable given that many resource strategies in insect larvae are unknown. Moreover, the species richness response may rely on the mobility and preferences of the observed imago rather than the occupation of the larvae. We therefore decided to follow a pragmatic division based on biological reasoning.

The response groups of our study included vascular plants, mosses, lichens, decomposing fungi, symbiotic fungi, flying insects (highly mobile insects dependent on ephemeral food sources such as dung, flowers, fungi and dead wood), herbivores (mobile insects dependent on live, sessile plants), detritivores (invertebrates dependent on dead carbon sources), predators (arthropods dependent on live animals). For details on grouping, see Table S3. In order to investigate the potential for generalization across species groups we pooled species into the larger groups of producers (vascular plants, mosses and lichens), consumers (fungi and arthropods) and total (producers and consumers). In addition, we used richness of OTUs (operational taxonomic units (5)) of eukaryotes and fungi from soil eDNA and arthropods from eDNA extracted from ethanol from Malaise traps. For details on collecting species and eDNA data see Brunbjerg, *et al.* (11).

##### eDNA datasets

The preparation of the fungal and eukaryote eDNA datasets have been published in Frøslev, *et al.* (44) and Fløjgaard, *et al.* (45) respectively. The insect DNA dataset was produced by extracting DNA from the ethanol from the bulk insect Malaise traps and metabarcoding with an insect specific 16S primer. 45 ml ethanol and 1.5 ml of 3M sodium acetate were added to a 50 ml centrifuge tube, and left in a freezer for DNA precipitation overnight, then centrifuged for 40 minutes. The dried pellet was extracted with the Qiagen DNeasy blood and tissue kit (Qiagen, Germany) with minor modifications. The extracted DNA was normalized, amplified, sequenced and analyzed according to the overall procedures described for 16S insects in Fløjgaard, *et al.* (45), but including curation of OTUs with LULU (26) and taxonomic assignment with a custom script, and exclusion of non-arthropod sequences. Data from the two different collecting events were handled separately and the sequences were then combined for each site. Bioinformatic processing including links to all data is documented at https://github.com/tobiasgf/biowide_synthesis.

### Explanatory variables and statistical analyses

We built generalized linear models (GLMs) to predict species richness of selected response groups based on the best selection of ecospace variables. In addition, we built GLMs to predict the summed richness of vascular plants, mosses and lichens (producers) and fungi and invertebrates (consumers). Finally, we built an overall species richness model predicting the total richness of all observed species.

For each model we made a preliminary screening and selection of relevant variables, excluding variables with no hypothesized relationship to the species group or variables dependent on the response variable. We further constrained the response direction and shape to ecologically plausible responses (46, 47) implying an exclusion of negative effects of expansion, continuity and heterogeneity variables on species richness – more resources, more diverse resources, more environmental variation and increasing temporal and spatial continuity are all hypothesized to increase species richness if anything. This decision is justified by the large number of variables in our study increasing the risk of including spurious correlations in the models and thereby covering important causal relationships (Table S4).

Log transformation was preferred if model improvement was indicated by Akaike’s Information Criterion (AIC) (48). The number of explanatory variables were further reduced in order to avoid collinearity (VIF values < 3, (47)). A preliminary set of full models was built using all remaining variables: a general linear mixed poisson model (GLMM) with region as random variable and a GLM with poisson errors using the log link function. We selected the best model type using the ΔAIC < 2 criterium (46). Negative binomial errors were used if overdispersion was detected (49) in poisson models. We included a quadratic term of the abiotic position variables if the full model significantly improved according to the ΔAIC < 2 criterium. Expansion and continuity variables having a negative effect in the full model after variable transformation and adding of quadratic terms were deleted sequentially starting with the variable with the lowest z-value. The residuals of full models were checked for model misfit, overdispersion and spatial autocorrelation using simulated residuals and R package DHARMa (50). We used backwards elimination of explanatory variables using the ΔAIC < 2 rule to reduce full models to final models. For the flying insect (p=0.018) and DNA flying insect models (p=0.006), significant autocorrelation (p = 0.018 and p=0.006 in a Moran’s I test, respectively) was detected in the GLM negative binomial model. To avoid effects of autocorrelation on model selection, we used non-parametric model selection (leave-one-out cross validation) for these models while applying the ΔAIC < 2 rule.

Model performance of the final models were evaluated using leave-one-out cross-validation. To evaluate the effect of generalizing we compared the performance of models on aggregated species groups with the performance of the corresponding models on the individual species groups. in order to do this pearson correlations between the sum of predictions from specific group models and the sum of observed species of producers, consumers and total (referred to as the summed predictions from nine species group models and summed predictions for producers and consumers), were calculated respectively.

Variation partitioning was calculated on final models for each component of ecospace (position, expansion, continuity and co-variables) as follows:

> *adjusted deviance explained (best model)*
>
> — – *adjusted deviance explained(model without target ecospace component)*

Data exploration was applied following Zuur, Ieno and Elphick (47). All analyses were performed in R version 3.5.0 (51).

## Acknowledgements

RE, IG, TL, US, LD, HHB, TGF, CF and AKB were supported by a grant from VILLUM FONDEN (Biowide, VKR-023343). JCS considers this work a contribution to his VILLUM Investigator project “Biodiversity Dynamics in a Changing World” funded by VILLUM FONDEN (grant 16549). We thank Greg Newman for field and laboratory support.

## Supplementary Information

Fig. S1: Nonlinear responses in a quasipoisson GAM of vascular plant optima frequencies in the Danish flora along Ellenberg factors representing variation in light, moisture and pH.

Table S2: Model estimates for the linear model used for calculating Soil Fertility Index.

Table S3: List of families belonging to the functional groups: lichens, decomposing fungi, symbiotic fungi, flying insects, herbivores, detritivores and predatory arthropods.

Table S4: Ecological plausible relationships between explanatory variables (co-variables, position, expansion and continuity) and response groups used to constrain the terms entering our multiple regression models.

