## Supplemental material for "Multi-taxon inventory reveals highly consistent biodiversity responses to ecospace variation"

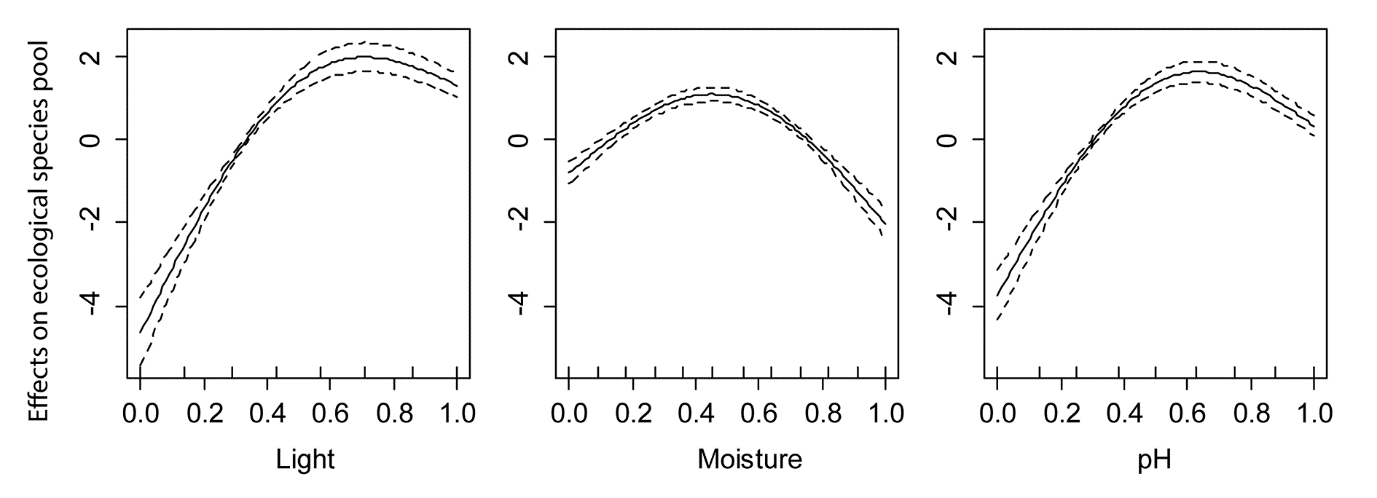

**Figure S1:** Nonlinear responses in a quasipoisson GAM of the preferences of vascular plants in the Danish flora along Ellenberg factors representing variation in light, moisture and pH. Most species prefer relatively high light and pH and intermediate soil moisture. The resulting predictions were log-transformed to produce the index for ecological species pool size.

**Table S2:** The index for soil fertility was derived by combining a number of indicators into an index predicting variation in Ellenberg N. Estimates, p-values (* <0.05, ** <0.01, *** <0.001) and standard errors (in parentheses) for the linear model: Ellenberg N ~ log(Ca^2+^ content) + leaf nitrogen + leaf nitrogen/phosphorus + soil class. The estimate for Soil = Clay is set to 0.000 by definition as this is the reference soil type for the categorical variable. N=520. R^2^=69.5%

|  | Ellenberg N |
| --- | --- |
| Intercept | 1.209^***^ (0.233) |
| Log (Ca^2+^ content) | 0.396^***^ (0.023) |
| Leaf nitrogen | 0.695^***^ (0.042) |
| Leaf nitrogen/phosphorus | -0.083^***^ (0.009) |
| Soil = Organic | -1.214^***^ (0.109) |
| Soil = Sand | -0.572^***^ (0.106) |
| Soil = Sand/clay | -0.420^***^ (0.110) |
| Soil = Clay | 0.000 |

**Table S3:** List of families and genera belonging to the functional groups: lichens, decomposing fungi (decomposers), symbiotic fungi (symbionts), flying insects, herbivores, detritivores and predatory arthropods. Functional group affiliation is coarse and follows tractable taxonomic levels given data available.

| Functional group | Order | Family | Genus | Species |
| --- | --- | --- | --- | --- |
| Decomposer | Agaricales | Agaricaceae | Agaricus |  |
| Decomposer | Agaricales | Agaricaceae | Bovista |  |
| Decomposer | Agaricales | Agaricaceae | Chlorophyllum |  |
| Decomposer | Agaricales | Agaricaceae | Coprinus |  |
| Decomposer | Agaricales | Agaricaceae | Cyathus |  |
| Decomposer | Agaricales | Agaricaceae | Cystoderma |  |
| Decomposer | Agaricales | Agaricaceae | Cystodermella |  |
| Decomposer | Agaricales | Agaricaceae | Cystolepiota |  |
| Decomposer | Agaricales | Agaricaceae | Disciseda |  |
| Decomposer | Agaricales | Agaricaceae | Lepiota |  |
| Decomposer | Agaricales | Agaricaceae | Leucoagaricus |  |
| Decomposer | Agaricales | Agaricaceae | Lycoperdon |  |
| Decomposer | Agaricales | Agaricaceae | Macrolepiota |  |
| Decomposer | Agaricales | Agaricaceae | Melanophyllum |  |
| Decomposer | Agaricales | Agaricaceae | Mycocalia |  |
| Decomposer | Boletales | Amylocorticiaceae | Amylocorticium |  |
| Decomposer | Boletales | Amylocorticiaceae | Amyloxenasma |  |
| Decomposer | Boletales | Amylocorticiaceae | Ceraceomyces |  |
| Decomposer | Russulales | Amylostereaceae | Amylostereum |  |
| Decomposer | Pezizales | Ascobolaceae | Ascobolus |  |
| Decomposer | Helotiales | Ascocorticiaceae | Ascocorticium |  |
| Decomposer | Pezizales | Ascodesmidaceae | Lasiobolus |  |
| Decomposer | Atheliales | Atheliaceae | Athelia |  |
| Decomposer | Atheliales | Atheliaceae | Athelopsis |  |
| Decomposer | Atheliales | Atheliaceae | Hypochniciellum |  |
| Decomposer | Atheliales | Atheliaceae | Leptosporomyces |  |
| Decomposer | Atheliales | Atheliaceae | Tretomyces |  |
| Decomposer | Auriculariales | Auriculariaceae | Auricularia |  |
| Decomposer | Auriculariales | Auriculariaceae | Eichleriella |  |
| Decomposer | Auriculariales | Auriculariaceae | Exidiopsis |  |
| Decomposer | Russulales | Auriscalpiaceae | Auriscalpium |  |
| Decomposer | Russulales | Auriscalpiaceae | Clavicorona |  |
| Decomposer | Russulales | Auriscalpiaceae | Lentinellus |  |
| Decomposer | Coronophorales | Bertiaceae | Bertia |  |
| Decomposer | Agaricales | Bolbitiaceae | Bolbitius |  |
| Decomposer | Agaricales | Bolbitiaceae | Conocybe |  |
| Decomposer | Agaricales | Bolbitiaceae | Pholiotina |  |
| Decomposer | Boliniales | Boliniaceae | Camarops |  |
| Decomposer | Cantharellales | Botryobasidiaceae | Botryobasidium |  |
| Decomposer | Cantharellales | Botryobasidiaceae | Botryohypochnus |  |
| Decomposer | Coronophorales | Chaetosphaerellaceae | Chaetosphaerella |  |
| Decomposer | Agaricales | Clavariaceae | Mucronella |  |
| Decomposer | Boletales | Coniophoraceae | Coniophora |  |
| Decomposer | Corticiales | Corticiaceae | Corticium |  |
| Decomposer | Corticiales | Corticiaceae | Dendrothele |  |
| Decomposer | Corticiales | Corticiaceae | Lyomyces |  |
| Decomposer | Agaricales | Cyphellaceae | Seticyphella |  |
| Decomposer | Dacrymycetales | Dacrymycetaceae | Calocera |  |
| Decomposer | Dacrymycetales | Dacrymycetaceae | Dacrymyces |  |
| Decomposer | Incertae sedis | Denticularia | Denticularia |  |
| Decomposer | Agaricales | Entolomataceae | Clitopilus |  |
| Decomposer | Agaricales | Entolomataceae | Rhodocybe |  |
| Decomposer | Polyporales | Fomitopsidaceae | Anomoporia |  |
| Decomposer | Polyporales | Fomitopsidaceae | Antrodia |  |
| Decomposer | Polyporales | Fomitopsidaceae | Dacryobolus |  |
| Decomposer | Polyporales | Fomitopsidaceae | Daedalea |  |
| Decomposer | Polyporales | Fomitopsidaceae | Fomitopsis |  |
| Decomposer | Polyporales | Fomitopsidaceae | Ischnoderma |  |
| Decomposer | Polyporales | Fomitopsidaceae | Oligoporus |  |
| Decomposer | Polyporales | Fomitopsidaceae | Postia |  |
| Decomposer | Geastrales | Geastraceae | Geastrum |  |
| Decomposer | Geastrales | Geastraceae | Sphaerobolus |  |
| Decomposer | Gloeophyllales | Gloeophyllaceae | Gloeophyllum |  |
| Decomposer | Gomphales | Gomphaceae | Ceratellopsis |  |
| Decomposer | Gomphales | Gomphaceae | Ramaria |  |
| Decomposer | Sordariales | Helminthosphaeriaceae | Echinosphaeria |  |
| Decomposer | Sordariales | Helminthosphaeriaceae | Ruzenia |  |
| Decomposer | Helotiales | Helotiaceae | Chlorociboria |  |
| Decomposer | Helotiales | Helotiaceae | Ombrophila |  |
| Decomposer | Helotiales | Helotiaceae | Unguiculariopsis |  |
| Decomposer | Russulales | Hericiaceae | Laxitextum |  |
| Decomposer | Chaetothyriales | Herpotrichiellaceae | Capronia |  |
| Decomposer | Tremellales | Hyaloriaceae | Myxarium |  |
| Decomposer | Helotiales | Hyaloscyphaceae | Arachnopeziza |  |
| Decomposer | Helotiales | Hyaloscyphaceae | Dematioscypha |  |
| Decomposer | Helotiales | Hyaloscyphaceae | Hyaloscypha |  |
| Decomposer | Helotiales | Hyaloscyphaceae | Lasiobelonium |  |
| Decomposer | Helotiales | Hyaloscyphaceae | Olla |  |
| Decomposer | Helotiales | Hyaloscyphaceae | Pseudolachnea |  |
| Decomposer | Cantharellales | Hydnaceae | Paullicorticium |  |
| Decomposer | Cantharellales | Hydnaceae | Sistotrema |  |
| Decomposer | Trechisporales | Hydnodontaceae | Brevicellicium |  |
| Decomposer | Trechisporales | Hydnodontaceae | Luellia |  |
| Decomposer | Trechisporales | Hydnodontaceae | Sistotremastrum |  |
| Decomposer | Trechisporales | Hydnodontaceae | Sistotremella |  |
| Decomposer | Trechisporales | Hydnodontaceae | Sphaerobasidium |  |
| Decomposer | Trechisporales | Hydnodontaceae | Subulicystidium |  |
| Decomposer | Trechisporales | Hydnodontaceae | Trechispora |  |
| Decomposer | Trechisporales | Hydnodontaceae | Tubulicium |  |
| Decomposer | Agaricales | Hygrophoraceae | Ampulloclitocybe |  |
| Decomposer | Boletales | Hygrophoropsidaceae | Hygrophoropsis |  |
| Decomposer | Boletales | Hygrophoropsidaceae | Leucogyrophana |  |
| Decomposer | Hymenochaetales | Hymenochaetaceae | Fuscoporia |  |
| Decomposer | Hymenochaetales | Hymenochaetaceae | Hymenochaete |  |
| Decomposer | Hymenochaetales | Hymenochaetaceae | Inonotus |  |
| Decomposer | Hymenochaetales | Hymenochaetaceae | Tubulicrinis |  |
| Decomposer | Incertae sedis | Incertae sedis | Basidiodendron |  |
| Decomposer | Incertae sedis | Incertae sedis | Chalara |  |
| Decomposer | Incertae sedis | Incertae sedis | Endoperplexa |  |
| Decomposer | Incertae sedis | Incertae sedis | Farlowiella |  |
| Decomposer | Incertae sedis | Incertae sedis | Hauerslevia |  |
| Decomposer | Incertae sedis | Incertae sedis | Natantiella |  |
| Decomposer | Incertae sedis | Incertae sedis | Oliveonia |  |
| Decomposer | Incertae sedis | Incertae sedis | Panaeolus |  |
| Decomposer | Incertae sedis | Incertae sedis | Phlebiella |  |
| Decomposer | Incertae sedis | Incertae sedis | Plicatura |  |
| Decomposer | Incertae sedis | Incertae sedis | Pseudochaete |  |
| Decomposer | Incertae sedis | Incertae sedis | Resinicium |  |
| Decomposer | Incertae sedis | Incertae sedis | Stypella |  |
| Decomposer | Incertae sedis | Incertae sedis | Trechinothus |  |
| Decomposer | Incertae sedis | Incertae sedis | Trichaptum |  |
| Decomposer | Incertae sedis | Incertae sedis | Typhrasa |  |
| Decomposer | Incertae sedis | Incertae sedis | Xylohypha |  |
| Decomposer | Agaricales | Inocybaceae | Crepidotus |  |
| Decomposer | Agaricales | Inocybaceae | Episphaeria |  |
| Decomposer | Agaricales | Inocybaceae | Flammulaster |  |
| Decomposer | Agaricales | Inocybaceae | Phaeomarasmius |  |
| Decomposer | Agaricales | Inocybaceae | Simocybe |  |
| Decomposer | Boletales | Jaapiaceae | Jaapia |  |
| Decomposer | Russulales | Lachnocladiaceae | Asterostroma |  |
| Decomposer | Russulales | Lachnocladiaceae | Scytinostroma |  |
| Decomposer | Russulales | Lachnocladiaceae | Vararia |  |
| Decomposer | Sordariales | Lasiosphaeriaceae | Cercophora |  |
| Decomposer | Sordariales | Lasiosphaeriaceae | Lasiosphaeria |  |
| Decomposer | Sordariales | Lasiosphaeriaceae | Lasiosphaeris |  |
| Decomposer | Sordariales | Lasiosphaeriaceae | Podospora |  |
| Decomposer | Gomphales | Lentariaceae | Kavinia |  |
| Decomposer | Gomphales | Lentariaceae | Lentaria |  |
| Decomposer | Agaricales | Lyophyllaceae | Lyophyllum |  |
| Decomposer | Agaricales | Lyophyllaceae | Rugosomyces |  |
| Decomposer | Agaricales | Marasmiaceae | Baeospora |  |
| Decomposer | Agaricales | Marasmiaceae | Calyptella |  |
| Decomposer | Agaricales | Marasmiaceae | Cephaloscypha |  |
| Decomposer | Agaricales | Marasmiaceae | Crinipellis |  |
| Decomposer | Agaricales | Marasmiaceae | Hydropus |  |
| Decomposer | Agaricales | Marasmiaceae | Macrocystidia |  |
| Decomposer | Agaricales | Marasmiaceae | Marasmius |  |
| Decomposer | Agaricales | Marasmiaceae | Megacollybia |  |
| Decomposer | Agaricales | Marasmiaceae | Rectipilus |  |
| Decomposer | Polyporales | Meripilaceae | Physisporinus |  |
| Decomposer | Polyporales | Meruliaceae | Bjerkandera |  |
| Decomposer | Polyporales | Meruliaceae | Bulbillomyces |  |
| Decomposer | Polyporales | Meruliaceae | Cabalodontia |  |
| Decomposer | Polyporales | Meruliaceae | Cerocorticium |  |
| Decomposer | Polyporales | Meruliaceae | Hyphoderma |  |
| Decomposer | Polyporales | Meruliaceae | Hypochnicium |  |
| Decomposer | Polyporales | Meruliaceae | Junghuhnia |  |
| Decomposer | Polyporales | Meruliaceae | Lyomyces |  |
| Decomposer | Polyporales | Meruliaceae | Mycoaciella |  |
| Decomposer | Polyporales | Meruliaceae | Phlebia |  |
| Decomposer | Polyporales | Meruliaceae | Scopuloides |  |
| Decomposer | Polyporales | Meruliaceae | Steccherinum |  |
| Decomposer | Agaricales | Mycenaceae | Hemimycena |  |
| Decomposer | Agaricales | Mycenaceae | Mycena |  |
| Decomposer | Agaricales | Mycenaceae | Panellus |  |
| Decomposer | Agaricales | Mycenaceae | Resinomycena |  |
| Decomposer | Agaricales | Mycenaceae | Roridomyces |  |
| Decomposer | Agaricales | Mycenaceae | Sarcomyxa |  |
| Decomposer | Agaricales | Niaceae | Flagelloscypha |  |
| Decomposer | Agaricales | Niaceae | Lachnella |  |
| Decomposer | Agaricales | Omphalotaceae | Gymnopus |  |
| Decomposer | Agaricales | Omphalotaceae | Marasmiellus |  |
| Decomposer | Agaricales | Omphalotaceae | Mycetinis |  |
| Decomposer | Agaricales | Omphalotaceae | Rhodocollybia |  |
| Decomposer | Onygenales | Onygenaceae | Onygena |  |
| Decomposer | Orbiliales | Orbiliaceae | Hyalorbilia |  |
| Decomposer | Orbiliales | Orbiliaceae | Orbilia |  |
| Decomposer | Incertae sedis | Parodiopsidaceae | Henningsomyces |  |
| Decomposer | Patellariales | Patellariaceae | Rhizodiscina |  |
| Decomposer | Boletales | Paxillaceae | Hydnomerulius |  |
| Decomposer | Russulales | Peniophoraceae | Gloiothele |  |
| Decomposer | Russulales | Peniophoraceae | Peniophora |  |
| Decomposer | Russulales | Peniophoraceae | Vesiculomyces |  |
| Decomposer | Pezizales | Pezizaceae | Iodophanus |  |
| Decomposer | Phaeotrichales | Phaeotrichaceae | Trichodelitschia |  |
| Decomposer | Phallales | Phallaceae | Mutinus |  |
| Decomposer | Phallales | Phallaceae | Phallus |  |
| Decomposer | Polyporales | Phanerochaetaceae | Antrodiella |  |
| Decomposer | Polyporales | Phanerochaetaceae | Ceriporia |  |
| Decomposer | Polyporales | Phanerochaetaceae | Ceriporiopsis |  |
| Decomposer | Polyporales | Phanerochaetaceae | Meruliopsis |  |
| Decomposer | Polyporales | Phanerochaetaceae | Phanerochaete |  |
| Decomposer | Polyporales | Phanerochaetaceae | Rhizochaete |  |
| Decomposer | Atractiellales | Phleogenaceae | Helicogloea |  |
| Decomposer | Atractiellales | Phleogenaceae | Phleogena |  |
| Decomposer | Agaricales | Physalacriaceae | Cylindrobasidium |  |
| Decomposer | Agaricales | Physalacriaceae | Flammulina |  |
| Decomposer | Agaricales | Physalacriaceae | Gloiocephala |  |
| Decomposer | Agaricales | Physalacriaceae | Rhizomarasmius |  |
| Decomposer | Agaricales | Physalacriaceae | Strobilurus |  |
| Decomposer | Platygloeales | Platygloeaceae | Achroomyces |  |
| Decomposer | Agaricales | Pleurotaceae | Hohenbuehelia |  |
| Decomposer | Agaricales | Pluteaceae | Pluteus |  |
| Decomposer | Polyporales | Polyporaceae | Cinereomyces |  |
| Decomposer | Polyporales | Polyporaceae | Daedaleopsis |  |
| Decomposer | Polyporales | Polyporaceae | Datronia |  |
| Decomposer | Polyporales | Polyporaceae | Hapalopilus |  |
| Decomposer | Polyporales | Polyporaceae | Neofavolus |  |
| Decomposer | Polyporales | Polyporaceae | Polyporus |  |
| Decomposer | Polyporales | Polyporaceae | Skeletocutis |  |
| Decomposer | Polyporales | Polyporaceae | Trametes |  |
| Decomposer | Polyporales | Polyporaceae | Tyromyces |  |
| Decomposer | Agaricales | Porotheleaceae | Porotheleum |  |
| Decomposer | Agaricales | Psathyrellaceae | Coprinellus |  |
| Decomposer | Agaricales | Psathyrellaceae | Coprinopsis |  |
| Decomposer | Agaricales | Psathyrellaceae | Lacrymaria |  |
| Decomposer | Agaricales | Psathyrellaceae | Parasola |  |
| Decomposer | Agaricales | Psathyrellaceae | Psathyrella |  |
| Decomposer | Agaricales | Pterulaceae | Aphanobasidium |  |
| Decomposer | Agaricales | Pterulaceae | Globulicium |  |
| Decomposer | Agaricales | Pterulaceae | Merulicium |  |
| Decomposer | Agaricales | Pterulaceae | Pterula |  |
| Decomposer | Pezizales | Pyronemataceae | Cheilymenia |  |
| Decomposer | Pezizales | Pyronemataceae | Pseudombrophila |  |
| Decomposer | Pezizales | Pyronemataceae | Scutellinia |  |
| Decomposer | Pezizales | Pyronemataceae | Spooneromyces |  |
| Decomposer | Hymenochaetales | Repetobasidiaceae | Sidera |  |
| Decomposer | Hymenochaetales | Rickenellaceae | Peniophorella |  |
| Decomposer | Russulales | Russulaceae | Boidinia |  |
| Decomposer | Atractiellales | Saccoblastiaceae | Saccoblastia |  |
| Decomposer | Pezizales | Sarcoscyphaceae | Sarcoscypha |  |
| Decomposer | Agaricales | Schizophyllaceae | Schizophyllum |  |
| Decomposer | Hymenochaetales | Schizoporaceae | Basidioradulum |  |
| Decomposer | Hymenochaetales | Schizoporaceae | Hyphodontia |  |
| Decomposer | Hymenochaetales | Schizoporaceae | Kneiffiella |  |
| Decomposer | Hymenochaetales | Schizoporaceae | Xylodon |  |
| Decomposer | Boletales | Serpulaceae | Serpula |  |
| Decomposer | Russulales | Stephanosporaceae | Cristinia |  |
| Decomposer | Russulales | Stephanosporaceae | Lindtneria |  |
| Decomposer | Russulales | Stereaceae | Acanthobasidium |  |
| Decomposer | Russulales | Stereaceae | Aleurodiscus |  |
| Decomposer | Russulales | Stereaceae | Gloeocystidiellum |  |
| Decomposer | Russulales | Stereaceae | Stereum |  |
| Decomposer | Agaricales | Strophariaceae | Agrocybe |  |
| Decomposer | Agaricales | Strophariaceae | Deconica |  |
| Decomposer | Agaricales | Strophariaceae | Flammula |  |
| Decomposer | Agaricales | Strophariaceae | Gymnopilus |  |
| Decomposer | Agaricales | Strophariaceae | Hypholoma |  |
| Decomposer | Agaricales | Strophariaceae | Kuehneromyces |  |
| Decomposer | Agaricales | Strophariaceae | Leratiomyces |  |
| Decomposer | Agaricales | Strophariaceae | Phaeonematoloma |  |
| Decomposer | Agaricales | Strophariaceae | Pholiota |  |
| Decomposer | Agaricales | Strophariaceae | Protostropharia |  |
| Decomposer | Agaricales | Strophariaceae | Psilocybe |  |
| Decomposer | Agaricales | Strophariaceae | Stropharia |  |
| Decomposer | Boletales | Tapinellaceae | Tapinella |  |
| Decomposer | Thelebolales | Thelebolaceae | Ascozonus |  |
| Decomposer | Thelebolales | Thelebolaceae | Coprotus |  |
| Decomposer | Agaricales | Tricholomataceae | Aspropaxillus |  |
| Decomposer | Agaricales | Tricholomataceae | Cellypha |  |
| Decomposer | Agaricales | Tricholomataceae | Clitocybe |  |
| Decomposer | Agaricales | Tricholomataceae | Delicatula |  |
| Decomposer | Agaricales | Tricholomataceae | *Hygroaster* |  |
| Decomposer | Agaricales | Tricholomataceae | Infundibulicybe |  |
| Decomposer | Agaricales | Tricholomataceae | Lepista |  |
| Decomposer | Agaricales | Tricholomataceae | Leucocybe |  |
| Decomposer | Agaricales | Tricholomataceae | Melanoleuca |  |
| Decomposer | Agaricales | Tricholomataceae | Mycenella |  |
| Decomposer | Agaricales | Tricholomataceae | Pseudobaeospora |  |
| Decomposer | Agaricales | Tricholomataceae | Pseudoclitocybe |  |
| Decomposer | Agaricales | Tricholomataceae | Pseudolasiobolus |  |
| Decomposer | Agaricales | Tricholomataceae | Resupinatus |  |
| Decomposer | Agaricales | Tricholomataceae | Ripartites |  |
| Decomposer | Agaricales | Tricholomataceae | Tricholomopsis |  |
| Decomposer | Agaricales | Tubariaceae | Tubaria |  |
| Decomposer | Agaricales | Typhulaceae | Macrotyphula |  |
| Decomposer | Polyporales | Xenasmataceae | Xenasma |  |
| Decomposer | Xylariales | Xylariaceae | Hypocopra |  |
| Decomposer | Xylariales | Xylariaceae | Nemania |  |
| Decomposer | Xylariales | Xylariaceae | Poronia |  |
| Decomposer | Xylariales | Xylariaceae | Rosellinia |  |
| Decomposer | Agaricales | Physalacriaceae | Armillaria | A. lutea |
| Decomposer | Agaricales | Tricholomataceae | Arrhenia | A. epichysium |
| Decomposer | Agaricales | Entolomataceae | Entoloma | E. byssisedum; E. dichroum; E. dysthaloides; E. euchroum |
| Decomposer | Agaricales | Cortinariaceae | Galerina | G. embolus; G. lacustris; G. marginata; G. sideroides; G. triscopa; G. uncialis |
| Decomposer | Polyporales | Ganodermataceae | Ganoderma | G. applanatum |
| Decomposer | Helotiales | Dermateaceae | Mollisia | M. cinerea s.l. |
| Decomposer | Incertae sedis | Incertae sedis | Oxyporus | O. corticola |
| Decomposer | Pezizales | Pezizaceae | Peziza | P. varia |
| Decomposer | Xylariales | Xylariaceae | Xylaria | X. hypoxylon |
| Detritivores | Psocoptera | Amphipsocidae |  |  |
| Detritivores | Coleoptera | Anthribidae |  |  |
| Detritivores | Psocoptera | Caeciliusidae |  |  |
| Detritivores | Psocoptera | Ectopsocidae |  |  |
| Detritivores | Coleoptera | Elateridae |  |  |
| Detritivores | Psocoptera | Elipsocidae |  |  |
| Detritivores | Coleoptera | Endomychidae |  |  |
| Detritivores | Coleoptera | Eucnemidae |  |  |
| Detritivores | Hymenoptera | Formicidae |  |  |
| Detritivores | Psocoptera | Lachesillidae |  |  |
| Detritivores | Coleoptera | Latridiidae |  |  |
| Detritivores | Coleoptera | Melandryidae |  |  |
| Detritivores | Psocoptera | Mesopsocidae |  |  |
| Detritivores | Coleoptera | Monotomidae |  |  |
| Detritivores | Coleoptera | Nitidulidae |  |  |
| Detritivores | Psocoptera | Peripsocidae |  |  |
| Detritivores | Psocoptera | Philotarsidae |  |  |
| Detritivores | Diptera | Phoridae |  |  |
| Detritivores | Psocoptera | Psocidae |  |  |
| Detritivores | Coleoptera | Ptinidae |  |  |
| Detritivores | Diptera | Ptychopteridae |  |  |
| Detritivores | Coleoptera | Pyrochroidae |  |  |
| Detritivores | Coleoptera | Salpingidae |  |  |
| Detritivores | Neuroptera | Sisyridae |  |  |
| Detritivores | Psocoptera | Stenopsocidae |  |  |
| Detritivores | Coleoptera | Tenebrionidae |  |  |
| Detritivores | Coleoptera | Throscidae |  |  |
| Detritivores | Psocoptera | Trogiidae |  |  |
| Flying insects | Diptera | Acroceridae |  |  |
| Flying insects | Hymenoptera | Ampulicidae |  |  |
| Flying insects | Hymenoptera | Andrenidae |  |  |
| Flying insects | Hymenoptera | Aphelinidae |  |  |
| Flying insects | Hymenoptera | Apidae |  |  |
| Flying insects | Hymenoptera | Argidae |  |  |
| Flying insects | Diptera | Asilidae |  |  |
| Flying insects | Trichoptera | Beraeidae |  |  |
| Flying insects | Hymenoptera | Bethylidae |  |  |
| Flying insects | Diptera | Bombyliidae |  |  |
| Flying insects | Hymenoptera | Braconidae |  |  |
| Flying insects | Coleoptera | Cantharidae |  |  |
| Flying insects | Coleoptera | Cerambycidae |  |  |
| Flying insects | Hymenoptera | Chalcididae |  |  |
| Flying insects | Hymenoptera | Chrysididae |  |  |
| Flying insects | Hymenoptera | Colletidae |  |  |
| Flying insects | Hymenoptera | Crabronidae |  |  |
| Flying insects | Lepidoptera | Crambidae |  |  |
| Flying insects | Hymenoptera | Diapriidae |  |  |
| Flying insects | Lepidoptera | Drepanidae |  |  |
| Flying insects | Hymenoptera | Dryinidae |  |  |
| Flying insects | Trichoptera | Ecnomidae |  |  |
| Flying insects | Lepidoptera | Elachistidae |  |  |
| Flying insects | Strepsiptera | Elenchidae |  |  |
| Flying insects | Hymenoptera | Encyrtidae |  |  |
| Flying insects | Lepidoptera | Erebidae |  |  |
| Flying insects | Hymenoptera | Eurytomidae |  |  |
| Flying insects | Hymenoptera | Evaniidae |  |  |
| Flying insects | Hymenoptera | Gasteruptiidae |  |  |
| Flying insects | Coleoptera | Geotrupidae |  |  |
| Flying insects | Hymenoptera | Halictidae |  |  |
| Flying insects | Strepsiptera | Halictophagidae |  |  |
| Flying insects | Lepidoptera | Hepialidae |  |  |
| Flying insects | Lepidoptera | Hesperiidae |  |  |
| Flying insects | Coleoptera | Histeridae |  |  |
| Flying insects | Diptera | Hybotidae |  |  |
| Flying insects | Coleoptera | Hydrophilidae |  |  |
| Flying insects | Trichoptera | Hydropsychidae |  |  |
| Flying insects | Trichoptera | Hydroptilidae |  |  |
| Flying insects | Hymenoptera | Ichneumonidae |  |  |
| Flying insects | Lepidoptera | Lasiocampidae |  |  |
| Flying insects | Trichoptera | Lepidostomatidae |  |  |
| Flying insects | Trichoptera | Leptoceridae |  |  |
| Flying insects | Lepidoptera | Limacodidae |  |  |
| Flying insects | Trichoptera | Limnephilidae |  |  |
| Flying insects | Coleoptera | Lucanidae |  |  |
| Flying insects | Lepidoptera | Lycaenidae |  |  |
| Flying insects | Hymenoptera | Megachilidae |  |  |
| Flying insects | Hymenoptera | Megaspilidae |  |  |
| Flying insects | Hymenoptera | Melittidae |  |  |
| Flying insects | Diptera | Micropezidae |  |  |
| Flying insects | Trichoptera | Molannidae |  |  |
| Flying insects | Hymenoptera | Mutillidae |  |  |
| Flying insects | Hymenoptera | Mymaridae |  |  |
| Flying insects | Lepidoptera | Nymphalidae |  |  |
| Flying insects | Coleoptera | Oedemeridae |  |  |
| Flying insects | Diptera | Oestridae |  |  |
| Flying insects | Diptera | Pediciidae |  |  |
| Flying insects | Trichoptera | Phryganeidae |  |  |
| Flying insects | Lepidoptera | Pieridae |  |  |
| Flying insects | Hymenoptera | Platygastridae |  |  |
| Flying insects | Diptera | Platystomatidae |  |  |
| Flying insects | Trichoptera | Polycentropodidae |  |  |
| Flying insects | Hymenoptera | Pompilidae |  |  |
| Flying insects | Trichoptera | Psychomyiidae |  |  |
| Flying insects | Hymenoptera | Pteromalidae |  |  |
| Flying insects | Diptera | Rhagionidae |  |  |
| Flying insects | Diptera | Rhinophoridae |  |  |
| Flying insects | Coleoptera | Scarabaeidae |  |  |
| Flying insects | Diptera | Scathophagidae |  |  |
| Flying insects | Diptera | Sciomyzidae |  |  |
| Flying insects | Trichoptera | Sericostomatidae |  |  |
| Flying insects | Coleoptera | Silphidae |  |  |
| Flying insects | Hymenoptera | Sphecidae |  |  |
| Flying insects | Lepidoptera | Sphingidae |  |  |
| Flying insects | Coleoptera | Staphylinidae |  |  |
| Flying insects | Diptera | Stratiomyidae |  |  |
| Flying insects | Diptera | Syrphidae |  |  |
| Flying insects | Diptera | Tachinidae |  |  |
| Flying insects | Plecoptera | Taeniopterygidae |  |  |
| Flying insects | Hymenoptera | Tiphiidae |  |  |
| Flying insects | Diptera | Ulidiidae |  |  |
| Flying insects | Hymenoptera | Vespidae |  |  |
| Flying insects | Diptera | Xylomyidae |  |  |
| Flying insects | Lepidoptera | Zygaenidae |  |  |
| Herbivores | Hemiptera | Acanthosomatidae |  |  |
| Herbivores | Orthoptera | Acrididae |  |  |
| Herbivores | Hemiptera | Alydidae |  |  |
| Herbivores | Hemiptera | Aphrophoridae |  |  |
| Herbivores | Coleoptera | Attelabidae |  |  |
| Herbivores | Hemiptera | Berytidae |  |  |
| Herbivores | Coleoptera | Brentidae |  |  |
| Herbivores | Coleoptera | Buprestidae |  |  |
| Herbivores | Hemiptera | Caliscelidae |  |  |
| Herbivores | Diptera | Cecidomyiidae |  |  |
| Herbivores | Hemiptera | Ceratocombidae |  |  |
| Herbivores | Hemiptera | Cercopidae |  |  |
| Herbivores | Diptera | Chloropidae |  |  |
| Herbivores | Hemiptera | Cicadellidae |  |  |
| Herbivores | Hemiptera | Cixiidae |  |  |
| Herbivores | Hemiptera | Coreidae |  |  |
| Herbivores | Coleoptera | Curculionidae |  |  |
| Herbivores | Hemiptera | Cydnidae |  |  |
| Herbivores | Hymenoptera | Cynipidae |  |  |
| Herbivores | Coleoptera | Dascillidae |  |  |
| Herbivores | Coleoptera | Dasytidae |  |  |
| Herbivores | Hemiptera | Delphacidae |  |  |
| Herbivores | Hemiptera | Dipsocoridae |  |  |
| Herbivores | Lepidoptera | Geometridae |  |  |
| Herbivores | Hemiptera | Hebridae |  |  |
| Herbivores | Lepidoptera | Incurvariidae |  |  |
| Herbivores | Coleoptera | Kateretidae |  |  |
| Herbivores | Coleoptera | Lycidae |  |  |
| Herbivores | Hemiptera | Lygaeidae |  |  |
| Herbivores | Coleoptera | Malachiidae |  |  |
| Herbivores | Hemiptera | Miridae |  |  |
| Herbivores | Lepidoptera | Noctuidae |  |  |
| Herbivores | Hymenoptera | Pamphiliidae |  |  |
| Herbivores | Hemiptera | Pemphigidae |  |  |
| Herbivores | Hemiptera | Pentatomidae |  |  |
| Herbivores | Orthoptera | Phaneropteridae |  |  |
| Herbivores | Hemiptera | Piesmatidae |  |  |
| Herbivores | Diptera | Psilidae |  |  |
| Herbivores | Hemiptera | Psyllidae |  |  |
| Herbivores | Lepidoptera | Pyralidae |  |  |
| Herbivores | Hemiptera | Rhopalidae |  |  |
| Herbivores | Hemiptera | Saldidae |  |  |
| Herbivores | Hemiptera | Scutelleridae |  |  |
| Herbivores | Hymenoptera | Tenthredinidae |  |  |
| Herbivores | Diptera | Tephritidae |  |  |
| Herbivores | Orthoptera | Tetrigidae |  |  |
| Herbivores | Orthoptera | Tettigoniidae |  |  |
| Herbivores | Hemiptera | Tingidae |  |  |
| Herbivores | Lepidoptera | Tortricidae |  |  |
| Herbivores | Hemiptera | Triozidae |  |  |
| Lichens | Acarosporales | Acarosporaceae | Acarospora |  |
| Lichens | Acarosporales | Acarosporaceae | Polysporina |  |
| Lichens | Arthoniales | Arthoniaceae | Arthonia |  |
| Lichens | Pleosporales | Arthopyreniaceae | Arthopyrenia |  |
| Lichens | Baeomycetales | Baeomycetaceae | Baeomyces |  |
| Lichens | Teloschistales | Caliciaceae | Amandinea |  |
| Lichens | Candelariales | Candelariaceae | Candelariella |  |
| Lichens | Lecanorales | Cladoniaceae | Cladonia |  |
| Lichens | Ostropales | Coenogoniaceae | Coenogonium |  |
| Lichens | Incertae sedis | Coniocybaceae | Chaenotheca |  |
| Lichens | Ostropales | Gomphillaceae | Jamesiella |  |
| Lichens | Ostropales | Graphidaceae | Graphis |  |
| Lichens | Ostropales | Graphidaceae | Thelotrema |  |
| Lichens | Ostropales | Gyalectaceae | Gyalecta |  |
| Lichens | Lecanorales | Haematommataceae | Haematomma |  |
| Lichens | Lecanorales | Lecanoraceae | Circinaria |  |
| Lichens | Lecanorales | Lecanoraceae | Lecanora |  |
| Lichens | Lecanorales | Lecanoraceae | Lecidella |  |
| Lichens | Lecanorales | Lecanoraceae | Protoparmeliopsis |  |
| Lichens | Lecanorales | Lecanoraceae | Pyrrhospora |  |
| Lichens | Lecanorales | Lecideaceae | Lecidea |  |
| Lichens | Lecanorales | Lecideaceae | Porpidia |  |
| Lichens | Pertusariales | Megasporaceae | Aspicilia |  |
| Lichens | Monoblastiales | Monoblastiaceae | Anisomeridium |  |
| Lichens | Lecanorales | Mycoblastaceae | Mycoblastus |  |
| Lichens | Mycocaliciales | Mycocaliciaceae | Mycocalicium |  |
| Lichens | Mycocaliciales | Mycocaliciaceae | Stenocybe |  |
| Lichens | Incertae sedis | Naetrocymbaceae | Leptorhaphis |  |
| Lichens | Pertusariales | Ochrolechiaceae | Ochrolechia |  |
| Lichens | Incertae sedis | Ophioparmaceae | Hypocenomyce |  |
| Lichens | Lecanorales | Parmeliaceae | Bryoria |  |
| Lichens | Lecanorales | Parmeliaceae | Cetraria |  |
| Lichens | Lecanorales | Parmeliaceae | Evernia |  |
| Lichens | Lecanorales | Parmeliaceae | Flavoparmelia |  |
| Lichens | Lecanorales | Parmeliaceae | Hypogymnia |  |
| Lichens | Lecanorales | Parmeliaceae | Hypotrachyna |  |
| Lichens | Lecanorales | Parmeliaceae | Melanelixia |  |
| Lichens | Lecanorales | Parmeliaceae | Melanohalea |  |
| Lichens | Lecanorales | Parmeliaceae | Parmelia |  |
| Lichens | Lecanorales | Parmeliaceae | Parmelina |  |
| Lichens | Lecanorales | Parmeliaceae | Parmeliopsis |  |
| Lichens | Lecanorales | Parmeliaceae | Parmotrema |  |
| Lichens | Lecanorales | Parmeliaceae | Platismatia |  |
| Lichens | Lecanorales | Parmeliaceae | Pleurosticta |  |
| Lichens | Lecanorales | Parmeliaceae | Pseudevernia |  |
| Lichens | Lecanorales | Parmeliaceae | Punctelia |  |
| Lichens | Lecanorales | Parmeliaceae | Tuckermannopsis |  |
| Lichens | Lecanorales | Parmeliaceae | Usnea |  |
| Lichens | Lecanorales | Parmeliaceae | Xanthoparmelia |  |
| Lichens | Peltigerales | Peltigeraceae | Peltigera |  |
| Lichens | Pertusariales | Pertusariaceae | Pertusaria |  |
| Lichens | Pertusariales | Pertusariaceae | Pseudosagedia |  |
| Lichens | Ostropales | Phlyctidaceae | Phlyctis |  |
| Lichens | Teloschistales | Physciaceae | Buellia |  |
| Lichens | Teloschistales | Physciaceae | Calicium |  |
| Lichens | Teloschistales | Physciaceae | Diplotomma |  |
| Lichens | Teloschistales | Physciaceae | Phaeophyscia |  |
| Lichens | Teloschistales | Physciaceae | Physcia |  |
| Lichens | Teloschistales | Physciaceae | Rinodina |  |
| Lichens | Lecanorales | Pilocarpaceae | Micarea |  |
| Lichens | Teloschistales | Psoraceae | Protoblastenia |  |
| Lichens | Lecanorales | Ramalinaceae | Bacidia |  |
| Lichens | Lecanorales | Ramalinaceae | Cliostomum |  |
| Lichens | Lecanorales | Ramalinaceae | Lecania |  |
| Lichens | Lecanorales | Ramalinaceae | Ramalina |  |
| Lichens | Incertae sedis | Rhizocarpaceae | Rhizocarpon |  |
| Lichens | Arthoniales | Roccellaceae | Alyxoria |  |
| Lichens | Arthoniales | Roccellaceae | Dendrographa |  |
| Lichens | Arthoniales | Roccellaceae | Lecanactis |  |
| Lichens | Arthoniales | Roccellaceae | Opegrapha |  |
| Lichens | Incertae sedis | Sarrameanaceae | Loxospora |  |
| Lichens | Lecanorales | Scoliciosporaceae | Scoliciosporum |  |
| Lichens | Lecanorales | Stereocaulaceae | Lepraria |  |
| Lichens | Teloschistales | Teloschistaceae | Athallia |  |
| Lichens | Teloschistales | Teloschistaceae | Calogaya |  |
| Lichens | Teloschistales | Teloschistaceae | Gyalolechia |  |
| Lichens | Teloschistales | Teloschistaceae | Polycauliona |  |
| Lichens | Teloschistales | Teloschistaceae | Xanthoria |  |
| Lichens | Baeomycetales | Trapeliaceae | Placynthiella |  |
| Lichens | Baeomycetales | Trapeliaceae | Trapelia |  |
| Lichens | Baeomycetales | Trapeliaceae | Trapeliopsis |  |
| Lichens | Verrucariales | Verrucariaceae | Pyrenula |  |
| Lichens | Verrucariales | Verrucariaceae | Verrucaria |  |
| Predatory arthropods | Araneae | Agelenidae |  |  |
| Predatory arthropods | Araneae | Amaurobiidae |  |  |
| Predatory arthropods | Hemiptera | Anthocoridae |  |  |
| Predatory arthropods | Araneae | Anyphaenidae |  |  |
| Predatory arthropods | Araneae | Araneidae |  |  |
| Predatory arthropods | Araneae | Atypidae |  |  |
| Predatory arthropods | Coleoptera | Carabidae |  |  |
| Predatory arthropods | Neuroptera | Chrysopidae |  |  |
| Predatory arthropods | Hemiptera | Cimicidae |  |  |
| Predatory arthropods | Coleoptera | Cleridae |  |  |
| Predatory arthropods | Araneae | Clubionidae |  |  |
| Predatory arthropods | Coleoptera | Coccinellidae |  |  |
| Predatory arthropods | Neuroptera | Coniopterygidae |  |  |
| Predatory arthropods | Araneae | Corinnidae |  |  |
| Predatory arthropods | Araneae | Cybaeidae |  |  |
| Predatory arthropods | Araneae | Dictynidae |  |  |
| Predatory arthropods | Coleoptera | Drilidae |  |  |
| Predatory arthropods | Prostigmata | Eriophyidae |  |  |
| Predatory arthropods | Prostigmata | Erythraeidae |  |  |
| Predatory arthropods | Hemiptera | Gerridae |  |  |
| Predatory arthropods | Araneae | Gnaphosidae |  |  |
| Predatory arthropods | Araneae | Hahniidae |  |  |
| Predatory arthropods | Neuroptera | Hemerobiidae |  |  |
| Predatory arthropods | Coleoptera | Lampyridae |  |  |
| Predatory arthropods | Araneae | Linyphiidae |  |  |
| Predatory arthropods | Araneae | Liocranidae |  |  |
| Predatory arthropods | Araneae | Lycosidae |  |  |
| Predatory arthropods | Hemiptera | Microphysidae |  |  |
| Predatory arthropods | Araneae | Mimetidae |  |  |
| Predatory arthropods | Araneae | Miturgidae |  |  |
| Predatory arthropods | Hemiptera | Nabidae |  |  |
| Predatory arthropods | Opiliones | Nemastomatidae |  |  |
| Predatory arthropods | Hemiptera | Nepidae |  |  |
| Predatory arthropods | Araneae | Oxyopidae |  |  |
| Predatory arthropods | Opiliones | Phalangiidae |  |  |
| Predatory arthropods | Araneae | Philodromidae |  |  |
| Predatory arthropods | Araneae | Pisauridae |  |  |
| Predatory arthropods | Raphidioptera | Raphidiidae |  |  |
| Predatory arthropods | Hemiptera | Reduviidae |  |  |
| Predatory arthropods | Araneae | Salticidae |  |  |
| Predatory arthropods | Araneae | Segestriidae |  |  |
| Predatory arthropods | Araneae | Sparassidae |  |  |
| Predatory arthropods | Araneae | Tetragnathidae |  |  |
| Predatory arthropods | Araneae | Theridiidae |  |  |
| Predatory arthropods | Araneae | Theridiosomatidae |  |  |
| Predatory arthropods | Araneae | Thomisidae |  |  |
| Predatory arthropods | Opiliones | Trogulidae |  |  |
| Predatory arthropods | Araneae | Uloboridae |  |  |
| Predatory arthropods | Hemiptera | Veliidae |  |  |
| Predatory arthropods | Araneae | Zoridae |  |  |
| Symbiont | Acrospermales | Acrospermaceae | Acrospermum |  |
| Symbiont | Agaricales | Agaricaceae | Amanita |  |
| Symbiont | Xylariales | Apiosporaceae | Arthrinium |  |
| Symbiont | Rhytismatales | Ascodichaenaceae | Ascodichaena |  |
| Symbiont | Atheliales | Atheliaceae | Amphinema |  |
| Symbiont | Atheliales | Atheliaceae | Byssocorticium |  |
| Symbiont | Atheliales | Atheliaceae | Piloderma |  |
| Symbiont | Atheliales | Atheliaceae | Tylospora |  |
| Symbiont | Auriculariales | Auriculariaceae | Exidia |  |
| Symbiont | Hypocreales | Bionectriaceae | Hydropisphaera |  |
| Symbiont | Hypocreales | Bionectriaceae | Nectriopsis |  |
| Symbiont | Boletales | Boletaceae | Boletus |  |
| Symbiont | Boletales | Boletaceae | Caloboletus |  |
| Symbiont | Boletales | Boletaceae | Chalciporus |  |
| Symbiont | Boletales | Boletaceae | Hortiboletus |  |
| Symbiont | Boletales | Boletaceae | Imleria |  |
| Symbiont | Boletales | Boletaceae | Leccinum |  |
| Symbiont | Boletales | Boletaceae | Neoboletus |  |
| Symbiont | Boletales | Boletaceae | Suillellus |  |
| Symbiont | Boletales | Boletaceae | Tylopilus |  |
| Symbiont | Boletales | Boletaceae | Xerocomellus |  |
| Symbiont | Boletales | Boletaceae | Xerocomus |  |
| Symbiont | Russulales | Bondarzewiaceae | Heterobasidion |  |
| Symbiont | Leotiales | Bulgariaceae | Bulgaria |  |
| Symbiont | Cantharellales | Cantharellaceae | Cantharellus |  |
| Symbiont | Cantharellales | Cantharellaceae | Craterellus |  |
| Symbiont | Cantharellales | Ceratobasidiaceae | Ceratobasidium |  |
| Symbiont | Cantharellales | Ceratobasidiaceae | Thanatephorus |  |
| Symbiont | Agaricales | Chromocyphellaceae | Chromocyphella |  |
| Symbiont | Agaricales | Clavariaceae | Capitoclavaria |  |
| Symbiont | Agaricales | Clavariaceae | Clavaria |  |
| Symbiont | Agaricales | Clavariaceae | Clavulinopsis |  |
| Symbiont | Agaricales | Clavariaceae | Hyphodontiella |  |
| Symbiont | Agaricales | Clavariaceae | Ramariopsis |  |
| Symbiont | Hypocreales | Clavicipitaceae | Claviceps |  |
| Symbiont | Cantharellales | Clavulinaceae | Clavulina |  |
| Symbiont | Hypocreales | Cordycipitaceae | Beauveria |  |
| Symbiont | Hypocreales | Cordycipitaceae | Cordyceps |  |
| Symbiont | Hypocreales | Cordycipitaceae | Gibellula |  |
| Symbiont | Hypocreales | Cordycipitaceae | Isaria |  |
| Symbiont | Hypocreales | Cordycipitaceae | Torrubiella |  |
| Symbiont | Corticiales | Corticiaceae | Vuilleminia |  |
| Symbiont | Agaricales | Cortinariaceae | Cortinarius |  |
| Symbiont | Agaricales | Cortinariaceae | Galerina |  |
| Symbiont | Agaricales | Cortinariaceae | Hebeloma |  |
| Symbiont | Pleosporales | Cucurbitariaceae | Cucurbitaria |  |
| Symbiont | Agaricales | Cyphellaceae | Chondrostereum |  |
| Symbiont | Helotiales | Dermateaceae | Cejpia |  |
| Symbiont | Helotiales | Dermateaceae | Diplonaevia |  |
| Symbiont | Helotiales | Dermateaceae | Mollisia |  |
| Symbiont | Helotiales | Dermateaceae | Niptera |  |
| Symbiont | Helotiales | Dermateaceae | Pezicula |  |
| Symbiont | Helotiales | Dermateaceae | Pseudopeziza |  |
| Symbiont | Helotiales | Dermateaceae | Trochila |  |
| Symbiont | Xylariales | Diatrypaceae | Diatrype |  |
| Symbiont | Xylariales | Diatrypaceae | Diatrypella |  |
| Symbiont | Xylariales | Diatrypaceae | Eutypa |  |
| Symbiont | Xylariales | Diatrypaceae | Eutypella |  |
| Symbiont | Xylariales | Diatrypaceae | Quaternaria |  |
| Symbiont | Dothideales | Dothideaceae | Phyllachora |  |
| Symbiont | Eurotiales | Elaphomycetaceae | Elaphomyces |  |
| Symbiont | Endogonales | Endogonaceae | Endogone |  |
| Symbiont | Agaricales | Entolomataceae | Entoloma |  |
| Symbiont | Platygloeales | Eocronartiaceae | Eocronartium |  |
| Symbiont | Exobasidiales | Exobasidiaceae | Exobasidium |  |
| Symbiont | Agaricales | Fistulinaceae | Fistulina |  |
| Symbiont | Polyporales | Fomitopsidaceae | Buglossoporus |  |
| Symbiont | Polyporales | Fomitopsidaceae | Laetiporus |  |
| Symbiont | Polyporales | Fomitopsidaceae | Piptoporus |  |
| Symbiont | Geoglossales | Geoglossaceae | Geoglossum |  |
| Symbiont | Geoglossales | Geoglossaceae | Glutinoglossum |  |
| Symbiont | Geoglossales | Geoglossaceae | Microglossum |  |
| Symbiont | Geoglossales | Geoglossaceae | Trichoglossum |  |
| Symbiont | Glomerales | Glomeraceae | Glomus |  |
| Symbiont | Helotiales | Helotiaceae | Ascocoryne |  |
| Symbiont | Helotiales | Helotiaceae | Ascotremella |  |
| Symbiont | Helotiales | Helotiaceae | Bisporella |  |
| Symbiont | Helotiales | Helotiaceae | Claussenomyces |  |
| Symbiont | Helotiales | Helotiaceae | Crocicreas |  |
| Symbiont | Helotiales | Helotiaceae | Cudoniella |  |
| Symbiont | Helotiales | Helotiaceae | Discinella |  |
| Symbiont | Helotiales | Helotiaceae | Godronia |  |
| Symbiont | Helotiales | Helotiaceae | Gorgoniceps |  |
| Symbiont | Helotiales | Helotiaceae | Hymenoscyphus |  |
| Symbiont | Helotiales | Helotiaceae | Ionomidotis |  |
| Symbiont | Helotiales | Helotiaceae | Phaeohelotium |  |
| Symbiont | Helotiales | Helotiaceae | Velutarina |  |
| Symbiont | Pezizales | Helvellaceae | Helvella |  |
| Symbiont | Helotiales | Hyaloscyphaceae | Calycellina |  |
| Symbiont | Helotiales | Hyaloscyphaceae | Calycina |  |
| Symbiont | Helotiales | Hyaloscyphaceae | Eriopezia |  |
| Symbiont | Helotiales | Hyaloscyphaceae | Incrucipulum |  |
| Symbiont | Helotiales | Hyaloscyphaceae | Lachnellula |  |
| Symbiont | Helotiales | *Hyaloscyphaceae* | Pezizella |  |
| Symbiont | Helotiales | Hyaloscyphaceae | Proliferodiscus |  |
| Symbiont | Helotiales | Hyaloscyphaceae | Psilachnum |  |
| Symbiont | Helotiales | Hyaloscyphaceae | Hamatocanthoscypha |  |
| Symbiont | Cantharellales | Hydnaceae | Hydnum |  |
| Symbiont | Agaricales | Hydnangiaceae | Laccaria |  |
| Symbiont | Agaricales | Hygrophoraceae | Cuphophyllus |  |
| Symbiont | Agaricales | Hygrophoraceae | Eonema |  |
| Symbiont | Agaricales | Hygrophoraceae | Gliophorus |  |
| Symbiont | Agaricales | Hygrophoraceae | Hygrocybe |  |
| Symbiont | Agaricales | Hygrophoraceae | Hygrophorus |  |
| Symbiont | Agaricales | Hygrophoraceae | Lichenomphalia |  |
| Symbiont | Hymenochaetales | Hymenochaetaceae | Fomitiporia |  |
| Symbiont | Hymenochaetales | Hymenochaetaceae | Mensularia |  |
| Symbiont | Hymenochaetales | Hymenochaetaceae | Phellinus |  |
| Symbiont | Hymenochaetales | Hymenochaetaceae | Phylloporia |  |
| Symbiont | Hymenochaetales | Hymenochaetaceae | Porodaedalea |  |
| Symbiont | Xylariales | Hyponectriaceae | Physalospora |  |
| Symbiont | Hysteriales | Hysteriaceae | Gloniopsis |  |
| Symbiont | Agaricales | Incertae sedis | Camarophyllopsis |  |
| Symbiont | Incertae sedis | Incertae sedis | Chaetospermum |  |
| Symbiont | Incertae sedis | Incertae sedis | Cotylidia |  |
| Symbiont | Agaricales | Incertae sedis | Gloioxanthomyces |  |
| Symbiont | Incertae sedis | Incertae sedis | Glyphium |  |
| Symbiont | Agaricales | Incertae sedis | Hodophilus |  |
| Symbiont | Incertae sedis | Incertae sedis | Homophron |  |
| Symbiont | Incertae sedis | Incertae sedis | Octavianina |  |
| Symbiont | Incertae sedis | Incertae sedis | Pseudotricholoma |  |
| Symbiont | Incertae sedis | Incertae sedis | Rhopographus |  |
| Symbiont | Hypocreales | Incertae sedis | Stilbella |  |
| Symbiont | Auriculariales | Incertae sedis | Tremellodendropsis |  |
| Symbiont | Agaricales | Inocybaceae | Inocybe |  |
| Symbiont | Helotiales | Lachnaceae | Capitotricha |  |
| Symbiont | Helotiales | Lachnaceae | Lachnum |  |
| Symbiont | Helotiales | *Lachnaceae* | Pezoloma |  |
| Symbiont | Helotiales | Lachnaceae | Trichopeziza |  |
| Symbiont | Helotiales | Leotiaceae | Leotia |  |
| Symbiont | Agaricales | Lyophyllaceae | Sphagnurus |  |
| Symbiont | Diaporthales | Melanconidaceae | Melanamphora |  |
| Symbiont | Diaporthales | Melanconidaceae | Melanconis |  |
| Symbiont | Pleosporales | Melanommataceae | Melanomma |  |
| Symbiont | Polyporales | Meripilaceae | Meripilus |  |
| Symbiont | Pezizales | Morchellaceae | Costantinella |  |
| Symbiont | Capnodiales | Mycosphaerellaceae | Cymadothea |  |
| Symbiont | Mytilinidiales | Mytilinidiaceae | Actidium |  |
| Symbiont | Mytilinidiales | Mytilinidiaceae | Lophium |  |
| Symbiont | Hypocreales | Nectriaceae | Cosmospora |  |
| Symbiont | Hypocreales | Nectriaceae | Nectria |  |
| Symbiont | Hypocreales | Nectriaceae | Neonectria |  |
| Symbiont | Hypocreales | Nectriaceae | Pleonectria |  |
| Symbiont | Agaricales | Niaceae | Maireina |  |
| Symbiont | Agaricales | Niaceae | Merismodes |  |
| Symbiont | Hypocreales | Ophiocordycipitaceae | Hirsutella |  |
| Symbiont | Hypocreales | Ophiocordycipitaceae | Ophiocordyceps |  |
| Symbiont | Boletales | Paxillaceae | Gyrodon |  |
| Symbiont | Boletales | Paxillaceae | Paxillus |  |
| Symbiont | Russulales | Peniophoraceae | Peniophora |  |
| Symbiont | Pezizales | Pezizaceae | Marcelleina |  |
| Symbiont | Pezizales | Pezizaceae | Pachyella |  |
| Symbiont | Pezizales | Pezizaceae | Peziza |  |
| Symbiont | Agaricales | Physalacriaceae | Armillaria |  |
| Symbiont | Agaricales | Physalacriaceae | Hymenopellis |  |
| Symbiont | Agaricales | Physalacriaceae | Mucidula |  |
| Symbiont | Agaricales | Physalacriaceae | Xerula |  |
| Symbiont | Polyporales | Polyporaceae | Aurantiporus |  |
| Symbiont | Polyporales | Polyporaceae | Epithele |  |
| Symbiont | Polyporales | Polyporaceae | Fomes |  |
| Symbiont | Pucciniales | Pucciniaceae | Gymnosporangium |  |
| Symbiont | Pucciniales | Pucciniaceae | Puccinia |  |
| Symbiont | Pucciniales | Pucciniastraceae | Pucciniastrum |  |
| Symbiont | Pezizales | Pyronemataceae | Geopora |  |
| Symbiont | Pezizales | Pyronemataceae | Humaria |  |
| Symbiont | Pezizales | Pyronemataceae | Leucoscypha |  |
| Symbiont | Pezizales | Pyronemataceae | Neottiella |  |
| Symbiont | Pezizales | Pyronemataceae | Octospora |  |
| Symbiont | Pezizales | Pyronemataceae | Otidea |  |
| Symbiont | Pezizales | Pyronemataceae | Parascutellinia |  |
| Symbiont | Pezizales | Pyronemataceae | Pulvinula |  |
| Symbiont | Pezizales | Pyronemataceae | Ramsbottomia |  |
| Symbiont | Pezizales | Pyronemataceae | Tarzetta |  |
| Symbiont | Pezizales | Pyronemataceae | Trichophaea |  |
| Symbiont | Rhytismatales | Rhytismataceae | Coccomyces |  |
| Symbiont | Rhytismatales | Rhytismataceae | Colpoma |  |
| Symbiont | Rhytismatales | Rhytismataceae | Hysterium |  |
| Symbiont | Rhytismatales | Rhytismataceae | Lophodermium |  |
| Symbiont | Rhytismatales | Rhytismataceae | Propolis |  |
| Symbiont | Rhytismatales | Rhytismataceae | Rhytisma |  |
| Symbiont | Rhytismatales | Rhytismataceae | Rhytismatales |  |
| Symbiont | Hymenochaetales | Rickenellaceae | Rickenella |  |
| Symbiont | Russulales | Russulaceae | Lactarius |  |
| Symbiont | Russulales | Russulaceae | Lactifluus |  |
| Symbiont | Russulales | Russulaceae | Russula |  |
| Symbiont | Helotiales | Rutstroemiaceae | Rutstroemia |  |
| Symbiont | Pezizales | Sarcoscyphaceae | Pithya |  |
| Symbiont | Boletales | Sclerodermataceae | Scleroderma |  |
| Symbiont | Helotiales | Sclerotiniaceae | Ciboria |  |
| Symbiont | Sebacinales | Sebacinaceae | Sebacina |  |
| Symbiont | Ostropales | Stictidaceae | Cryptodiscus |  |
| Symbiont | Ostropales | Stictidaceae | Schizoxylon |  |
| Symbiont | Ostropales | Stictidaceae | Stictis |  |
| Symbiont | Agaricales | Strophariaceae | Hymenogaster |  |
| Symbiont | Agaricales | Strophariaceae | Naucoria |  |
| Symbiont | Boletales | Suillaceae | Suillus |  |
| Symbiont | Taphrinales | Taphrinaceae | Taphrina |  |
| Symbiont | Thelephorales | Thelephoraceae | Amaurodon |  |
| Symbiont | Thelephorales | Thelephoraceae | Pseudotomentella |  |
| Symbiont | Thelephorales | Thelephoraceae | Thelephora |  |
| Symbiont | Thelephorales | Thelephoraceae | Tomentella |  |
| Symbiont | Thelephorales | Thelephoraceae | Tomentellopsis |  |
| Symbiont | Pleosporales | Thyridariaceae | Thyridaria |  |
| Symbiont | Agaricales | Tricholomataceae | Arrhenia |  |
| Symbiont | Agaricales | Tricholomataceae | Dermoloma |  |
| Symbiont | Agaricales | Tricholomataceae | Fayodia |  |
| Symbiont | Agaricales | Tricholomataceae | Rimbachia |  |
| Symbiont | Agaricales | Tricholomataceae | Tricholoma |  |
| Symbiont | Tubeufiales | Tubeufiaceae | Tubeufia |  |
| Symbiont | Cantharellales | Tulasnellaceae | Tulasnella |  |
| Symbiont | Agaricales | Typhulaceae | Pistillina |  |
| Symbiont | Agaricales | Typhulaceae | Typhula |  |
| Symbiont | Xylariales | Xylariaceae | Annulohypoxylon |  |
| Symbiont | Xylariales | Xylariaceae | Entoleuca |  |
| Symbiont | Xylariales | Xylariaceae | Euepixylon |  |
| Symbiont | Xylariales | Xylariaceae | Hypoxylon |  |
| Symbiont | Xylariales | Xylariaceae | Kretzschmaria |  |
| Symbiont | Xylariales | Xylariaceae | Lopadostoma |  |
| Symbiont | Xylariales | Xylariaceae | Xylaria |  |
| Symbiont | Atheliales | Atheliaceae | Athelia | A. arachnoidea |
| Symbiont | Polyporales | Ganodermataceae | Ganoderma | G. pfeifferi |
| Symbiont | Incertae sedis | Incertae sedis | Oxyporus | O. populinus |
| Symbiont | Agaricales | Strophariaceae | Pholiota | P. squarrosa |
| Symbiont | Cantharellales | Hydnaceae | Sistotrema | S. muscicola |
| Symbiont | Trechisporales | Hydnodontaceae | Trechispora | T. fastidiosa |

**Table S4:** Assumed relationships between explanatory variables (co-variables, position, expansion and continuity) and response groups. Green (+): positive relationship only; orange (±): either positive or negative relationship; red (NA): variable not relevant for or circular with the response group. PA = presence/absence.

|  |  | Plants | Mosses | Lichens | Producers | Symbionts | Decomposers | Detritivores | Flying insects | Predatory arthropods | Herbivores | Consumers | Total | eDNA fungi | DNA_flying insects | eDNA eukaryotes |
| --- | --- | --- | --- | --- | --- | --- | --- | --- | --- | --- | --- | --- | --- | --- | --- | --- |
| Position | Ecological species pool | + | NA | NA | + | NA | NA | NA | NA | NA | NA | NA | NA | NA | NA | NA |
|  | Soil pH | ± | ± | ± | ± | ± | ± | ± | ± | ± | ± | ± | ± | ± | ± | ± |
|  | Soil fertility | ± | ± | ± | ± | ± | ± | ± | ± | ± | ± | ± | ± | ± | ± | ± |
|  | Soil moisture | ± | ± | ± | ± | ± | ± | ± | ± | ± | ± | ± | ± | ± | ± | ± |
|  | Light intensity | ± | ± | ± | ± | ± | ± | ± | ± | ± | ± | ± | ± | ± | ± | ± |
|  | Air temperature | ± | ± | ± | ± | ± | ± | ± | ± | ± | ± | ± | ± | ± | ± | ± |
|  | Boulder (PA) | NA | ± | ± | ± | NA | NA | NA | NA | NA | NA | NA | ± | NA | NA | ± |
| Expansion | Litter mass | ± | ± | ± | ± | + | + | + | + | + | + | + | + | + | + | + |
|  | Soil organic C | ± | ± | ± | ± | + | + | + | + | + | + | + | + | + | + | + |
|  | Soil organic matter | ± | ± | ± | ± | + | + | + | + | + | + | + | + | + | + | + |
|  | Plant richness | NA | NA | NA | NA | + | + | + | + | + | + | + | NA | + | + | + |
|  | Flower abundance | NA | NA | NA | NA | NA | NA | NA | + | + | + | + | + | NA | + | + |
|  | Dung (PA) | NA | NA | NA | NA | + | + | + | + | + | NA | + | + | + | + | + |
|  | Dead wood volume | NA | + | + | + | + | + | + | + | + | + | + | + | + | + | + |
|  | Dead wood debris | NA | + | + | + | + | + | + | + | + | + | + | + | + | + | + |
|  | Fungi symbiont plants | NA | NA | NA | NA | + | + | NA | NA | NA | NA | + | NA | + | NA | + |
|  | Insect host plant availability | NA | NA | NA | NA | NA | NA | NA | + | + | + | + | NA | NA | + | + |
|  | Tree layer | NA | + | + | + | + | + | + | + | + | + | + | + | + | + | + |
|  | Shrub layer | NA | + | + | + | + | + | + | + | + | + | + | + | + | + | + |
|  | Density of large trees | NA | + | + | + | + | + | + | + | + | + | + | + | + | + | + |
| Continuity | Geological species pool | + | NA | NA | + | NA | NA | NA | NA | NA | NA | NA | NA | NA | NA | NA |
|  | Spatial continuity (500m) | + | + | + | + | + | + | + | + | + | + | + | + | + | + | + |
|  | Temporal continuity | + | + | + | + | + | + | + | + | + | + | + | + | + | + | + |
| Co-variables | Natural landscapes | ± | ± | ± | ± | ± | ± | ± | ± | ± | ± | ± | ± | ± | ± | ± |
|  | Soil pH variability | + | + | + | + | + | + | + | + | + | + | + | + | + | + | + |
|  | Soil moisture variability | + | + | + | + | + | + | + | + | + | + | + | + | + | + | + |
|  | Soil fertility variability | + | + | + | + | + | + | + | + | + | + | + | + | + | + | + |
|  | Tree layer variability | + | + | + | + | + | + | + | + | + | + | + | + | + | + | + |
|  | Shrub layer variability | + | + | + | + | + | + | + | + | + | + | + | + | + | + | + |
